# Ammonia inhibits energy metabolism in astrocytes in a rapid and GDH2-dependent manner

**DOI:** 10.1101/683763

**Authors:** Leonie Drews, Marcel Zimmermann, Rebecca E. Poss, Dominik Brilhaus, Laura Bergmann, Constanze Wiek, Roland P. Piekorz, Andreas P.M. Weber, Tabea Mettler-Altmann, Andreas S. Reichert

## Abstract

In hepatic encephalopathy (HE) astrocyte dysfunction is a primary factor impairing neuronal activity under hyperammonemia. We show that mitochondria in cellular HE models undergo rapid fragmentation under hyperammonemia in a reversible manner. Mitochondrial respiration and glycolysis were instantaneously hampered in a pH-independent manner. A metabolomics approach revealed a subsequent accumulation of numerous amino acids, including branched chain amino acids, and glucose. N^15^ labeling of ammonia shows rapid incorporation of ammonia-derived nitrogen into glutamate and glutamate-derived amino acids. Downregulating human *GLUD2*, encoding mitochondrial glutamate dehydrogenase 2 (GDH2), inhibiting GDH2 activity by SIRT4 overexpression, and supplementing cells with glutamate or glutamine alleviated ammonia-induced inhibition of mitochondrial respiration. Thus, under hyperammonemic conditions, GDH2 catalyzes the removal of ammonia by reductive amination of α-ketoglutarate but at the same time inhibits the TCA-cycle by depleting α-ketoglutarate. Overall, we propose a mitochondria-dependent mechanism contributing to the early steps in the pathogenesis of HE where the interplay between energy metabolism and ammonia removal plays a pivotal role.

## Introduction

Hepatic encephalopathy (HE) is a severe neuropsychiatric disorder caused by hyperammonemia due to various forms of acute or chronic liver dysfunction, most commonly liver cirrhosis. Another major cause of HE is portosystemic shunting, leading to the distribution of portal blood without removal of toxins in the liver (Cash et al., 2010). It is estimated that 50 to 70% of all patients suffering from liver cirrhosis develop minimal HE or HE (Al Sibae and McGuire, 2009; Patidar and Bajaj, 2015). The symptoms can vary from mild to severe, ranging from confusion or sleeping disturbances to hepatic coma and death (Ferenci et al., 2002). So far, the only curative treatment is liver transplantation (Larsen et al., 1995). Treatment options to reduce symptoms are manifold, but insufficient and have often not been subject to randomized control studies. The standard treatment includes the application of non-absorbable disaccharides, antibiotics or metabolic ammonia scavengers in order to reduce ammonia produced by gut-resident bacteria (Ferenci, 2017). A major precipitation factor in HE is hyperammonemia resulting from the impaired ability of the liver to eliminate ammonia via the urea cycle (Ferenci, 2017). Increased amounts of ammonia pass the blood-brain-barrier (Lockwood et al., 1980; Lockwood et al., 1979) and cause swelling and production of reactive oxygen and nitrogen species (ROS/RNS) in astrocytes (Görg et al., 2013; Norenberg et al., 2005). An increase of free radicals can lead to mitochondrial dysfunction, energy failure as well as formation of the mitochondrial transition pore (Albrecht and Norenberg, 2006; Bai et al., 2001, 2001; Frank et al., 2012; Stewart et al., 2000) or RNA oxidation (Görg et al., 2008). Several studies demonstrated that hyperammonemia primarily disturbs astrocyte function resulting in subsequent neurological dysfunction (Görg et al., 2018; Häussinger and Görg, 2010; Norenberg, 1987). Astrocytes have an essential role to ensure neuronal function, such as provision with nutrients and recycling of the neurotransmitter glutamate (Sontheimer, 1995; Zalc, 1994). Astrocytes are further involved in the detoxification of ammonia in the brain which is thought to occur primarily via ammonia fixation by glutamine synthetase in astrocytes (Martinez-Hernandez et al., 1977). However, the molecular mechanism of ammonia-induced neurological impairment is not clear. Ammonia detoxification in astrocytes by glutamine synthetase leads to the accumulation of glutamine, which causes astrocyte swelling (Cash et al., 2010) and can thereby lead to brain edema.

It has been suggested that ammonia can bind to the GABA receptor complex on astrocytes, which can lead to the production of neurosteroids. GABA is the main inhibitory neurotransmitter in the human brain and functions through the GABA receptor complex. Therefore, elevation in neurosteroids can inhibit neurotransmission (Ahboucha and Butterworth, 2004). Two neurosteroid precursors, allopregnanolone and pregnanolone, were found to be increased pathophysiologically in the brains of hepatic coma patients (Butterworth, 2016). Consistent with this, an increase of GABA has been shown in the cerebrospinal fluids (CSF) of HE patients (Al Sibae and McGuire, 2009). A recent analysis of the metabolome of CSF of HE patients additionally points towards alterations of metabolic pathways linked to energy metabolism (Weiss et al., 2016).

An increase in mitochondrial fission and hence in number of separated mitochondria was reported to occur in mice with severe liver damage in the *substantia nigra*, but not in the prefrontal cortex (Bai et al., 2018). In different HE rat models it was shown that the activity of respiratory chain complexes were decreased in different brain regions (Boer et al., 2009; Dhanda et al., 2017). Additionally, an impact on the TCA-cycle under hyperammonemia has been discussed, given that some TCA-cycle enzymes, *e.g.* pyruvate dehydrogenase and isocitrate dehydrogenase, were reported to be inhibited by ammonia (Katunuma et al., 1966; Zwingmann et al., 2003). The role of the TCA-cycle in the pathogenesis of HE is controversially discussed as *e.g.* conflicting results were found regarding the levels of different TCA-cycle intermediates (α-ketoglutarate, citrate and oxaloacetate) as well as the activity of α-ketoglutarate dehydrogenase (α-KGDH). Possible reasons for these discrepancies could be different HE models or the varying duration of ammonia exposure (reviewed in Rama Rao and Norenberg, 2012). Overall, along with disturbances of the mitochondrial morphology an imbalance of various energy metabolism pathways have been suggested as consequences of hyperammonemia, however, the effects and their possible contribution to the pathogenesis of HE are still unclear. In most studies prolonged treatments, such as 24 hours and beyond, have been used and are quite well studied (Görg et al., 2015; Hazell and Norenberg, 1998; Oenarto et al., 2016). However, the early events upon exposure to high ammonia concentrations are unclear. Therefore, we focused on effects after short durations of ammonia treatment and immediate outcomes thereof.

Disturbances of mitochondrial quality control mechanisms are known to be associated with ageing and numerous diseases, such as Optic atrophy, Parkinson’s disease, and many others (Orth and Schapira, 2002; Ramonet et al., 2013). The mitochondrial network constantly undergoes fission and fusion events, which are important to maintain mitochondrial quality control (Nunnari et al., 1997; Schäfer and Reichert, 2009). It was shown that different stress conditions, such as heat shock or increased ROS, can trigger fission of mitochondria leading to a fragmented mitochondrial morphology (Duvezin-Caubet et al., 2006; Frank et al., 2012). Mitochondria that are damaged lose their ability to fuse with the network and are removed by mitophagy, a selective form of autophagy (Frank et al., 2012; Twig et al., 2008). This mechanism prevents the spread of damage through the network and is an important measure to maintain a healthy mitochondrial network in the cell (Figge et al., 2013; Figge et al., 2012; Schäfer and Reichert, 2009). Polletta and colleagues have shown the appearance of small, round mitochondria in MDA-MB-231 human breast cancer and C2C12 mouse myoblast cells after treatment with NH_4_Cl among other stressors (Polletta et al., 2015).

Overall, it is not well understood whether and how altered mitochondrial function and energy metabolism could contribute to the pathogenesis of HE. In particular, it is not clear what happens at early time points after ammonia levels rise and what are later events occurring in astrocytes. Here we decided to address the effect of hyperammonemia on mitochondrial function and energy metabolism using a cellular model of HE. We show that ammonia impairs mitochondrial oxidative phosphorylation (OXPHOS) very rapidly, namely at a time scale of minutes. Moreover, we provide several lines of evidence that ammonia is primarily fixed by reductive amination of α-ketoglutarate to generate glutamate catalyzed by the mitochondrial glutamate dehydrogenase 2 (GDH2). This provides a novel view on the early steps of ammonia-induced toxicity and will help to better understand the pathogenesis of HE in future studies.

## Materials and methods

### Cell lines

Human astrocytoma cells (MOG-G-CCM) were derived from ECACC (European Collection of Authenticated Cell Cultures, Public Health England, Salisbury, UK) and are established from an anaplastic astrocytoma of human adult brain. Cells were grown in DMEM with 1 g/l Glucose (#D5546, Sigma-Aldrich, Taufkirchen, Germany) with 10% FCS (PAN-Biotech, Aidenbach, Germany), 2 mM GlutaMax (Thermo Fisher Scientific, Carlsbad CA, USA), 2 mM Sodium Pyruvate (Thermo Fisher Scientific, Carlsbad CA, USA) and Penicillin/Streptomycin (Merck Millipore, Burlington MA, USA) at 37 °C and 5% humidified CO_2_. Primary rat astrocytes were prepared from cerebral hemispheres of newborn Wistar rats and grown under the same conditions. A SIRT4-eGFP overexpressing HeLa cell line and respective eGFP control cell line were established as described below and grown under the conditions described above.

### Cellular metabolism analysis

Cells were plated according to the manufacturer’s instructions in Seahorse cell culture plate (Agilent Technologies, Santa Clara CA, USA) in respective growth media. Cells were seeded at a density resulting in a regular monolayer prior to analysis using a Seahorse XFe96 Extracellular Flux Analyzer (Agilent Technologies, Santa Clara CA, USA). Treatment with NH_4_Cl (VWR, Radnor PA, USA) or CH_3_NH_3_Cl (Merck Millipore, Burlington MA, USA) at the given molarity (5 mM if not indicated differently) and time was performed for 1, 4, 6, 24, 48 h, or immediately prior analysis (0 h) as indicated. One hour before analysis growth medium was removed, cells were washed with assay medium (#D5030, Sigma-Aldrich, Taufkirchen, Germany), and left on assay medium for measurement. The latter medium contained all supplements, lacked FCS and antibiotics, and contained NH_4_Cl or other compounds as indicated. For recovery tests, NH_4_Cl was omitted in the assay medium immediately prior to analysis. Reagents and protocols for Mito Stress and Glycolysis Stress Kits (Agilent Technologies, Santa Clara CA, USA) were used as indicated by the manufacturer. Cell plates were incubated at 37 °C in a non-CO_2_ incubator for 1 h. After measurement with the Seahorse XFe96 Analyzer (Agilent Technologies, Santa Clara CA, USA) cell numbers were quantified by absorption spectrometry (Ex.: 361 nm, Em.: 486 nm) using Hoechst 33342 (Thermo Fisher Scientific, Carlsbad CA, USA) staining using a plate reader (Tecan Infinite 200 PRO, Switzerland) and normalized to cell number. For treatment of cells with NH_4_Cl without preincubation (0 h), NH_4_Cl was added directly before the Seahorse plate was inserted into the machine after the non-CO_2_ incubation period. For rescue assays, cells were treated with respective compounds (10 mM) directly before treatment with 5 mM NH_4_Cl and the Mito Stress test was performed as described above. Analysis of cells transfected with siRNAs or plasmids occurred after 48 hours.

‘Maximal respiration’ is defined as the oxygen consumption rate (OCR) measured after Carbonyl-cyanide-4-(trifluoromethoxy)-phenylhydrazone (FCCP) injection (maximal OCR) minus the OCR measured after blocking mitochondrial respiration (non-mitochondrial OCR). Spare respiratory capacity is defined as ‘maximal OCR’ minus the initial OCR measured (basal respiration). Spare respiratory capacity in % is calculated as ‘maximal respiration’ / ‘basal respiration’ x 100. Maximal respiration and spare respiratory capacity are challenged states of respiration.

‘Glycolysis’ is defined as extracellular acidification rate (ECAR) and calculated as maximum rate measurement before oligomycin injection minus last rate of measurement before glucose injection. ‘Glycolytic Capacity’ is the maximum rate measurement after injection of oligomycin minus the last rate measurement before glucose injection.

### Plasmids and transfection of cell lines

To visualize mitochondria, the construct pEGFP-Mito (Clontech Laboratories, Mountain View CA, USA) was used. Transfection was done using Effectene Transfection Reagent (Qiagen, Hilden, Germany) according to the manufacturer’s instructions 24 h before imaging. Knock-down of glutamate dehydrogenase 2 (GDH2, encoded by the *GLUD2* gene) using siRNAs (#NM_012084: 5’CUAACCUCUUCACGUGUAA’3 and 5’UUACACGUGAAGAGGUUAG‘3, Sigma-Aldrich, Taufkirchen, Germany) or transfection of *GLUD2* plasmid/empty vector controls was done using Lipofectamine RNAiMAX (Thermo Fisher Scientific, Carlsbad CA, USA), according to the manufacturer’s instructions using the reverse transfection protocol. Analysis occurred 48 h after transfection. For *GLUD2* and empty vector plasmid transfections, as well as simultaneous transfection of plasmids and GDH siRNA, transfection was done directly in Seahorse cell plates. Human *GLUD2* cloned into pcDNA3.1(+), Clone ID: OHu18663, was obtained from GenScript USA Inc., Piscataway NJ, USA. As control vector the empty pcDNA3.1(+) vector (Invitrogen, Thermo Fisher Scientific, Carlsbad CA, USA) was used.

The cDNAs for eGFP and the human SIRT4-eGFP fusion protein were generated by PCR using pEGFP-N1 and pcDNA3.1-SIRT4-eGFP (Lang et al., 2017) as templates, respectively, and subsequently cloned via NheI and XhoI restriction sites into puc2CL12IPwo derived from plasmids generated earlier (Roellecke et al., 2016; Wiek et al., 2015). Constructs were verified by Sanger DNA sequencing. HEK293T cells were transfected as described (Roellecke et al., 2016; Wiek et al., 2015) using polyethylenimine transfection reagent (Sigma-Aldrich, Taufkirchen, Germany) with HIV1 helper plasmid (pCD/NL-BH) (Zhang et al., 2004), envelope vector (pczVSV-G) (Pietschmann et al., 1999), and the newly generated plasmids puc2CL12eGFPIPwo or puc2CL12SIRT4-eGFPIPwo (both containing an IRES-PuroR cassette). Viral supernatants were harvested 48 h after transfection, filtered through 0.45 µm filters (Sartorius AG, Göttingen, Germany), and used to transduce HeLa cells. Selection with 2 µg/ml Puromycin (InvivoGen, San Diego CA, USA) was started 96 h after transduction and eGFP positivity was tracked by flow cytometry (BD FACSCanto II, BD Biosciences, Franklin Lakes NJ, USA) in the FITC-A channel using non-transduced HeLa cells as negative control.

### Microscopy

For microscopic analysis, human astrocytoma cells were seeded in 3 cm glass bottom microscopy dishes (MatTek Corporation, Ashland MA, USA) and transfected with pEGFP-Mito. Cells were treated with 5 mM NH_4_Cl for periods of 1, 4, 6, 24, 48, or 72 h. Imaging was done with a Zeiss Axiovert Observer D1 microscope with 63x/1.4 NA oil objective (Filter: Ex 450-490 nm, Em 500-550 nm) (Zeiss, Oberkochen, Germany) and AxioVision Software. About 20 images per dish were taken and cells were categorized and quantitatively scored according to the degree of highly fragmented, fragmented, or tubular mitochondria. For analysis of reversibility of the mitochondrial phenotype, primary rat astrocytes transfected with pEGFP-Mito were first grown in the presence of 5 mM NH_4_Cl for 72 h and thereafter analyzed for mitochondrial fragmentation. Thereafter, medium was exchanged and cells were analyzed for mitochondrial morphology 24 h, 48 h, and 72 h after removal of NH_4_Cl. To determine mitochondrial morphology changes under the same conditions in primary rat astrocytes, mitochondria were visualized by immunostaining against Tom20 (primary antibody: sc-11415, Santa Cruz Biotechnology, Dallas TX, USA; secondary antibody: Alexa Fluor 488 #A27034, Thermo Fisher Scientific, Carlsbad CA, USA). Analysis was done as described above. Images were taken using a PerkinElmer Spinning Disk Confocal microscope (PerkinElmer, Waltham MA, USA) using the 488 nm laser, 60x/1.4 NA oil objective, and Volocity software. Images were processed with Fiji (Schindelin et al., 2012).

### Metabolite analysis

Cells were grown in 175 cm^2^ flasks (Greiner Bio-One, Kremsmünster, Austria) and treated with 5 mM NH_4_Cl for 48, 24, 6, 4, 2 or 1 h. For harvesting, cells were trypsinized (Trypsin: Sigma-Aldrich, Taufkirchen, Germany), washed twice with ice-cold PBS (Sigma-Aldrich, Taufkirchen, Germany) and counted. 1 x 10^6^ cells were pelleted and resuspended in a 1:2.5:1 mixture of H_2_O: methanol: chloroform (both VWR, Radnor PA, USA) pre-cooled to −20 °C. The suspension was mixed on a rotator at 4 °C for 10 min, centrifuged at 10,000 rpm for 5 min at 4 °C, and the supernatant was used for metabolite profiling as follows. (i) Polar metabolites were analyzed by gas chromatography coupled to a time-of-flight mass spectrometer (7200 GC-QTOF; Agilent Technologies, Santa Clara, CA, USA) according to Fiehn and Kind (Fiehn and Kind, 2007). For relative quantification, peak areas of the compounds were normalized to the peak area of the internal standard ribitol (Sigma-Aldrich, Taufkirchen, Germany) added to the extraction buffer. (ii) To follow accumulation of ^15^N label, cells were treated with ^15^NH_4_Cl as above. Amino acids were then analyzed by liquid chromatography coupled to a TOF MS (1290 UHPLC 6530 QTOF; Agilent Technologies, Santa Clara CA, USA) according to Gu et al. (2007). Relative ^15^N label enrichment was calculated after accounting for natural isotopic distribution. Peak integration and analysis were done using the Agilent Mass Hunter Workstation B07 (Agilent Technologies, Santa Clara CA, USA). The number of biological replicates for each experiment and further details are given in Supplemental Tables 1 to 3.

### Western Blot

Human astrocytoma cells were seeded in 6 cm dishes and transfected with siRNA specific for the GDH2 gene *GLUD2* as described above. Cells were harvested with 500 µl RIPA-buffer (Tris-HCl, NaCl, Triton X-100, sodium deoxycholate, SDS, EDTA, pH 7.4) with Complete protease inhibitor (Roche Diagnostics, Basel, Switzerland) and protein concentration was measured using a Bradford assay (Sigma-Aldrich, Taufkirchen, Germany). Proteins were separated with a 10% SDS-PAGE, blotted on a nitrocellulose membrane (Amersham, VWR, Radnor PA, USA), and blocked for 1 h with 5% fat-free milk powder solution (Carl Roth GmbH & Co. KG, Karlsruhe, Germany) in TBS. Primary antibody against GDH2 (Santa Cruz Biotechnology, Dallas TX, USA, sc-293459) was used at a 1:250 dilution over night; HSP60 antibody (Sigma-Aldrich, Taufkirchen, Germany, SAB4501464) as loading control was used at a 1:10,000 dilution for 1 h at RT. Blots were developed using Signal Fire ECL Reagent (Cell Signaling Technology, Danvers MA, USA) and visualized using the Fusion SL Gel Documentation System (PEQLAB, Germany).

### qPCR for GLUD knock-down validation

Human astrocytoma cells were seeded and transfected as described before. RNA extraction with All Prep RNA/Protein Kit (Qiagen) according to the manufacturer’s instructions. RNA was reverse transcribed to cDNA with QuantiNova Reverse Transcription Kit (Qiagen) according to the manufacturer’s instructions. 10 ng cDNA was used for qPCR with Rotorgene 6000 system (Corbett Research, now Qiagen) with the QuantiNova SYBR Green PCR Kit (Qiagen) according to the manufacturer’s instructions. Analysis was performed using Rotor-Gene Q Series Software (Qiagen) employing the ΔΔCt method. HPRT1 was used as housekeeping gene. qPCR primer: *GLUD2* forward: 5’-cggcagagttccaagacagt-3’; *GLUD2* reverse: 5’-gaacgctccattgtgtatgc-3’; HPRT1 forward: 5’-cctggcgtcgtgattagtg-3’; HPRT1 reverse: 5’-tgaggaataaacaccctttcca-3’.

### Statistics

Statistics was done using GraphPad Prism 7.04 for Windows (GraphPad Software, La Jolla CA, USA). For multiple comparisons One-way ANOVA with Dunnett’s or Tukey’s *post hoc* test was performed. To compare two groups Student’s t-test was applied. Data are shown ± SEM or SD as indicated.

## Results

### Ammonia induces mitochondrial fragmentation in a rapid and reversible manner

In order to test whether modulation of mitochondrial morphology by ammonia represents an early event in ammonia-induced effects in astrocytes, we first used the human astrocytoma cell line MOG-G-CCM transfected with pEGFP-Mito, a variant of GFP targeted to the mitochondrial matrix (Weber et al., 2013). The cells were treated with 5 mM NH_4_Cl for 1 to 72 hours and changes in mitochondrial morphology were quantified using confocal fluorescence microscopy. These or similar conditions are used in numerous established in vitro HE models (Görg et al., 2015; Warskulat et al., 2002) and also similar concentration of ammonia (5.4 mM) were present in the brain using an established in vivo rat model of HE (Swain et al., 1992). The appearance of cells showing enhanced mild (intermediate) or severe (highly fragmented) mitochondrial fragmentation (Fig. 1A) was evident already after 6 hours of treatment when compared to controls lacking NH_4_Cl or to earlier time points (Fig. 1BC). Conversely, the extent of mitochondria showing a tubular interconnected network-like morphology was markedly reduced by addition of ammonia. Mitochondrial fragmentation became more prominent after 24 h reaching a high steady-state level. Treatment for 48 h and 72 h with NH_4_Cl did not appear to further increase mitochondrial fragmentation (Fig. 1C). To corroborate this in primary cells we used an established in vitro HE model, namely primary rat astrocytes treated with 5 mM NH_4_Cl. Also here mitochondrial morphology (visualized by immunostaining against Tom20) was rapidly altered by NH_4_Cl (Fig. 1B). The change towards mitochondrial fragmentation was even more rapid as it already became evident after 1 h of ammonia treatment (Fig. 1D). Hence, mitochondrial morphology is altered very rapidly by hyperammonemia and primary rat astrocytes appear to even react faster than human astrocytoma cells. Next, we asked whether mitochondrial fragmentation is reversible upon removal of ammonia. Indeed, we observed in primary rat astrocytes that mitochondrial morphology, which was highly fragmented after 72 h treatment with NH_4_Cl, recovered within 24 h to a highly tubular morphology when the medium was exchanged to fresh medium lacking NH_4_Cl (Fig. S2). We conclude that mitochondrial fragmentation is rapidly induced by hyperammonemia within 1 to 6 h and is nearly fully reversible within 24 h after removal of ammonia.

**Figure 1:**
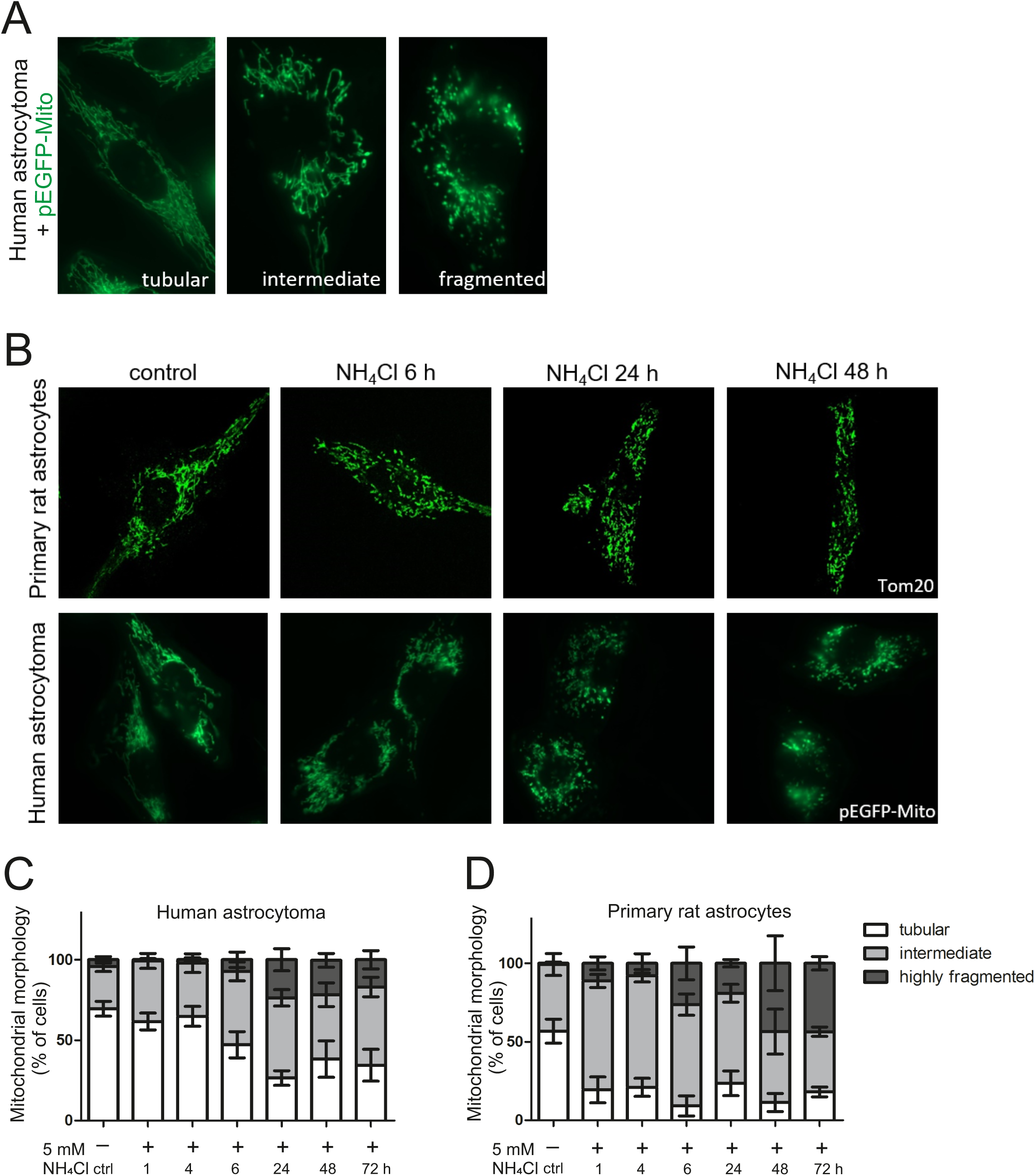
Mitochondrial morphology is altered by ammonia. Human astrocytoma cells were transfected with pEGFP-Mito to visualize mitochondria. In primary rat astrocytes visualization was achieved by immunostaining against Tom20. Cells were treated with 5 mM NH_4_Cl for indicated durations. At least 20 pictures were taken per sample showing approximately 3-5 cells each. Mitochondria were categorized into a tubular, intermediate, and fragmented morphological phenotype. (A) Representative images of morphological phenotype characterization in human astrocytoma cells. (B) Changes in morphology of mitochondria after treatment with 5 mM NH_4_Cl in primary rat astrocytes (top) and human astrocytoma (bottom). Time course of changes in mitochondrial morphology in human astrocytoma (C) and primary rat astrocytes (D). Percentage of cells with respective phenotype over time are shown. Data presented as mean ± SD (n=3-6).

### Ammonia causes an immediate inhibition of mitochondrial respiration in a pH-independent manner

Mitochondrial fragmentation is one of the early hallmarks indicating mitochondrial dysfunction (Duvezin-Caubet et al., 2006; Frank et al., 2012; Ishihara et al., 2006; Jheng et al., 2012). To investigate whether the observed rapid change in mitochondrial morphology comes hand in hand with mitochondrial dysfunction, we determined the cellular oxygen consumption rate (OCR) in human astrocytoma cells as well as in primary rat astrocytes after treatment with 5 mM NH_4_Cl for variable time periods. We applied the Mito Stress Kit using a Seahorse XFe96 Extracellular Flux Analyzer to determine different parameters of mitochondrial respiration including spare respiratory capacity and maximal respiration (for explanation see Fig. 2A and ‘Material and method’ section). Of note, these two parameters represent the mitochondrial respiration in a challenged state by uncoupling using FCCP. In human astrocytoma cells the spare respiratory capacity (Fig. 2B) as well as the maximal respiration (Fig. S1A) was significantly reduced already after 1 h of ammonia treatment compared to non-ammonia treated controls. Prolonged ammonia pretreatments up to 48 h likewise showed a significant impairment of mitochondrial respiration. We consistently observed the trend that respiration is more affected with short times of ammonia pretreatment compared to longer times although this effect is statistically not significant. To elaborate this further and to check how fast ammonia can affect mitochondrial respiration, we decided to include a measurement that determines the OCR immediately after 5 mM NH_4_Cl was added. Indeed, this was sufficient to grossly impair mitochondrial respiration as indicated by the fact that the spare respiratory capacity (Fig. 2C) as well as the maximal respiration (Fig. S1B) in primary rat astrocytes was significantly reduced. This reduction in respiration without pretreatment (0 h) was even stronger when compared to 1 h, 4 h or 6 h ammonia pretreatment conditions, demonstrating that ammonia has an immediate negative effect on mitochondrial respiration. We next tested whether this immediate effect of ammonia is dose-dependent. This is indeed the case as concentrations as low as 1 mM NH_4_Cl are sufficient to induce a substantial drop in respiration which becomes more prominent with higher concentrations of NH_4_Cl (Fig. 2D and Fig. S1C). Ammonia is a potent base causing alkalization of extra- and intracellular compartments even under buffering conditions. To analyze whether the observed effect is simply caused by a pH-mediated effect, we repeated the OCR experiments using 5 mM CH_3_NH_3_Cl, which cannot be metabolized but acts as a pH-mimetic compound to NH_4_Cl. This pH-mimetic did not impair mitochondrial respiration to any significant extent independent of the duration of pretreatment, except maximal respiration after 48 h (Fig. 2E and Fig. S1D). This clearly excludes that the observed effect of ammonia is due to putative changes in the pH. Moreover, we tested whether removal of ammonia for 1 h is sufficient to restore mitochondrial respiration in human astrocytoma cells that have been pretreated with ammonia for 1 to 48 h. One hour in medium without ammonia led to a full recovery of mitochondrial respiration independent of the duration of pretreatment (Fig. S3). This is consistent with our data showing that mitochondrial fragmentation is restored after removal of ammonia from the culture medium.

**Figure 2:**
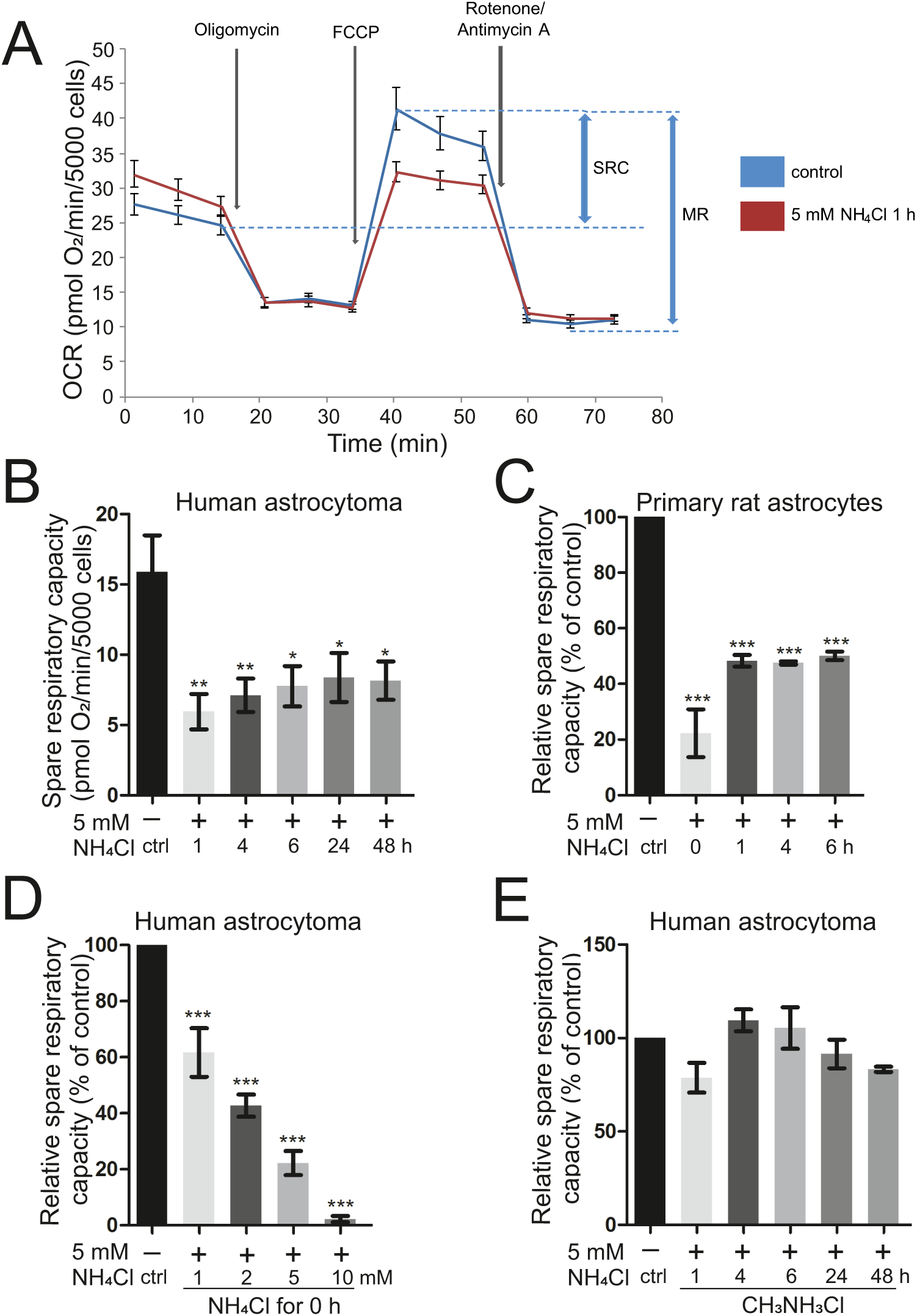
Mitochondrial respiration is immediately impaired by ammonia in a pH-independent manner. Oxygen consumption rate (OCR) of human astrocytoma cells and primary rat astrocytes was analyzed in Seahorse XFe96 Extracellular Flux Analyzer with the Mito Stress Test Kit after treatment with ammonia at the indicated molarities and time periods. (A) Scheme of Seahorse Mito Stress Test with indicated injections. SRC: spare respiratory capacity, MR: maximal respiration. (B) Spare respiratory capacity of human astrocytoma cells after treatment with 5 mM NH_4_Cl for 1-48 h (n=5-7). (C) Relative spare respiratory capacity of primary rat astrocytes after treatment with 5 mM NH_4_Cl for 1-6 h and directly after treatment (0 h) (n=3-4). (D) Relative spare respiratory capacity of human astrocytoma cells directly after treatment (0 h) with 1, 2, 5, or 10 mM NH_4_Cl (n=3). (E) Relative spare respiratory capacity of human astrocytoma cells after 5 mM CH_3_NH_3_Cl (pH-mimetic) treatment for 1-48 h (n=3). (C), (D), (E) Individual biological replicates normalized to controls (100%). Data represented as mean ± SEM. Statistics: One-way ANOVA with Dunnett’s post test. *P < 0.05, **P < 0.01, ***P < 0.001.

### Ammonia impairs glycolysis in astrocytes in a rapid manner

We examined the potential influence of ammonia on glycolytic flux and glycolytic capacity using the Glycolysis Stress Test with the Seahorse XFe96 Analyzer. Here the extracellular acidification rate (ECAR) is measured under different conditions and used to estimate glycolysis (see Fig. 3A and ‘Material and methods’ for explanations). Treatment of human astrocytoma cells with ammonia led to a significant and rapid reduction of glycolytic flux (Fig. 3B) and glycolytic capacity (Fig. 3C) when compared to controls. This was largely independent of the duration of ammonia pretreatment. Treatment with the pH-mimetic compound CH_3_NH_3_Cl did not have any detrimental effect on glycolysis further corroborating the specific role of ammonia independent from its property to alter the pH. Overall, we conclude that ammonia results in a very rapid, pH-independent, impairment of two major bioenergetic metabolic pathways, namely OXPHOS and glycolysis.

**Figure 3:**
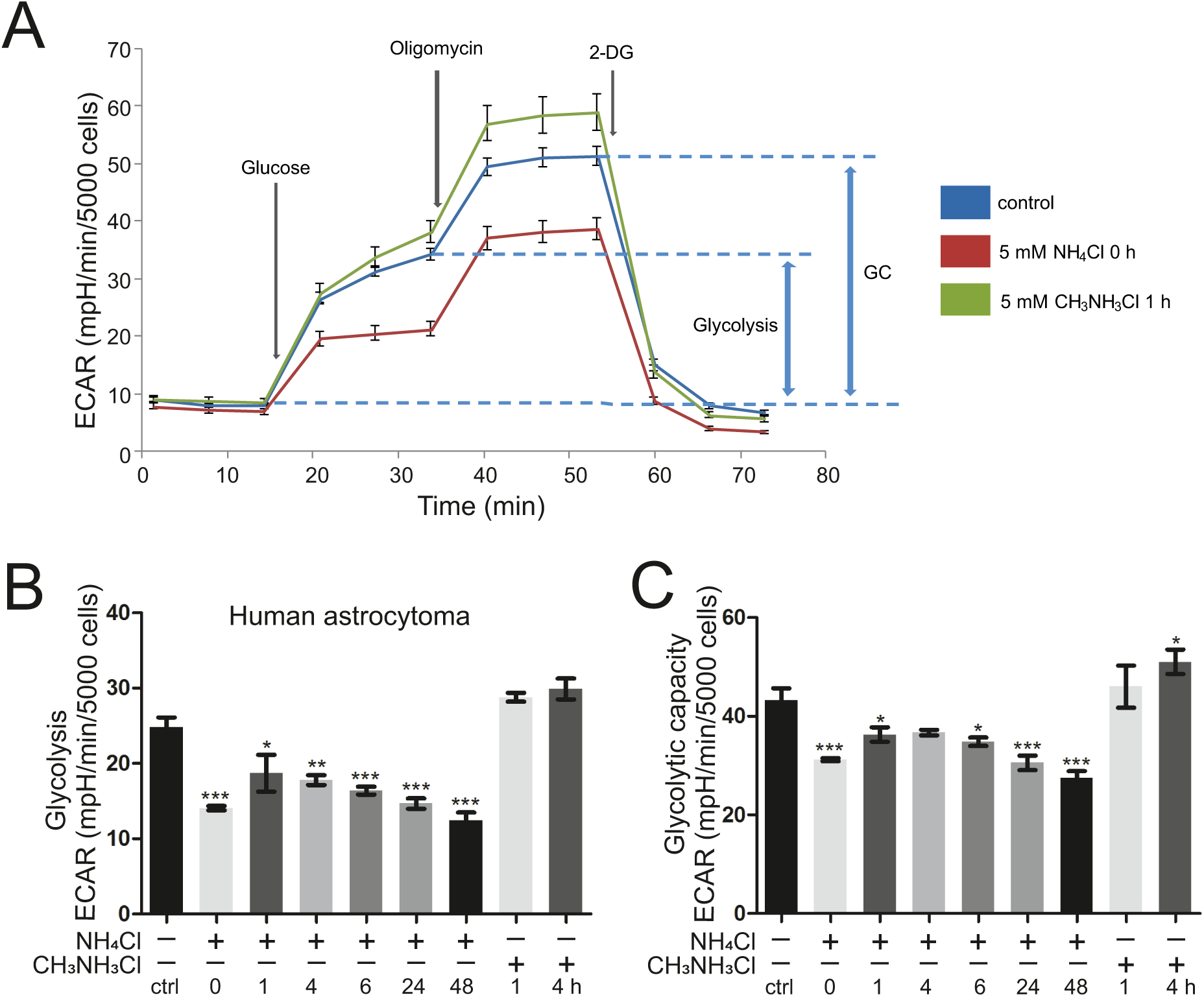
Ammonia reduces glycolytic function. Extracellular acidification rate (ECAR) was measured in human astrocytoma cells as a proxy for glycolysis using the Glycolysis Stress Test Kit on the Seahorse XFe96 Extracellular Flux Analyzer. (A) Scheme of the Seahorse Glycolysis Stress Test with indicated injections. GC: glycolytic capacity. (B) Glycolysis and (C) glycolytic capacity of human astrocytoma cells after treatment with 5 mM NH_4_Cl for 1-48 h and directly after treatment (0 h) (n=3); treatment with 5 mM pH-mimetic CH_3_NH_3_Cl for 1 or 4 h (n=2). Data represented as mean ± SEM. Statistics: One-way ANOVA with Dunnett’s post test. *P < 0.05, **P < 0.01, ***P < 0.001.

**Figure 4:**
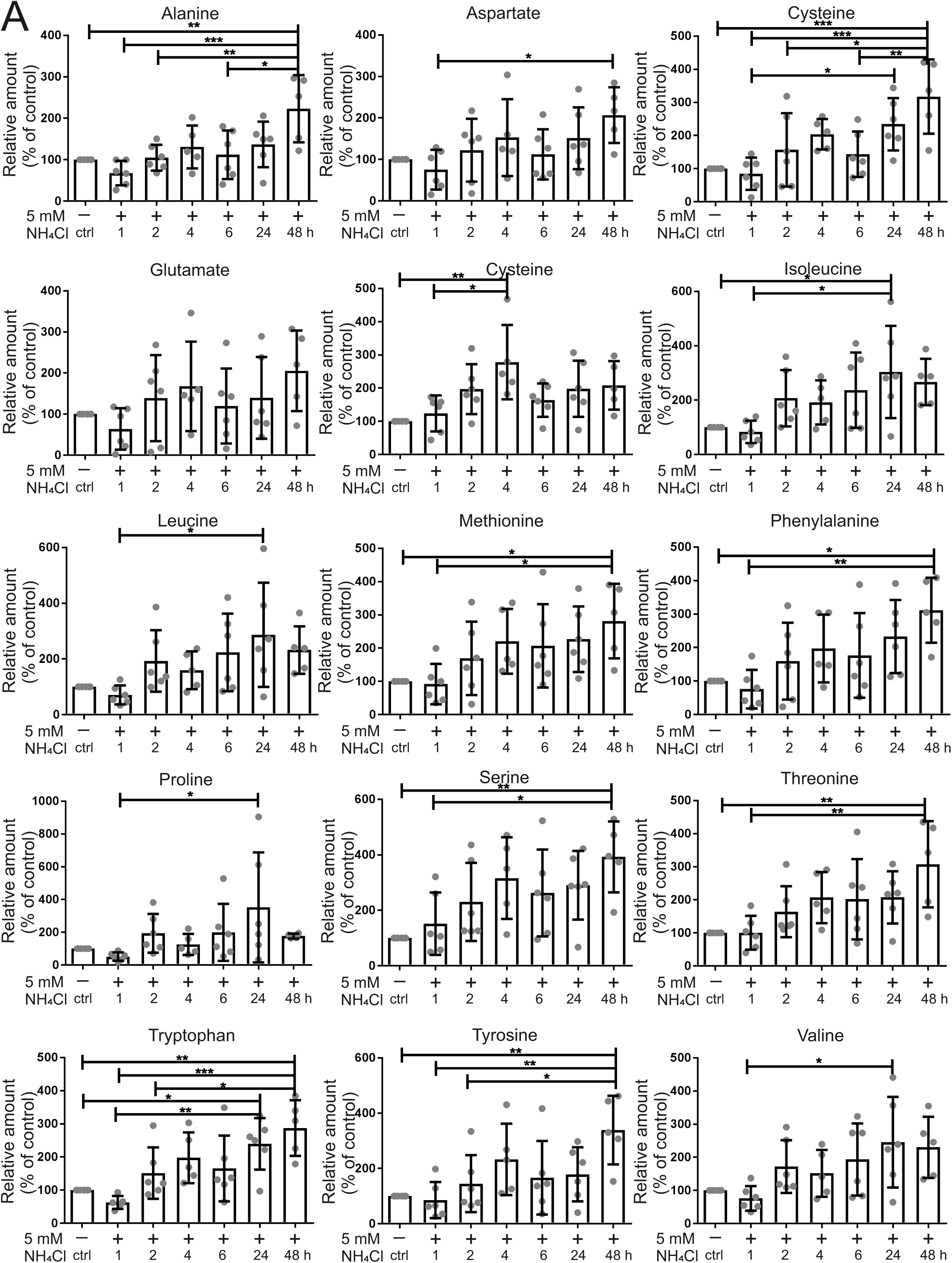
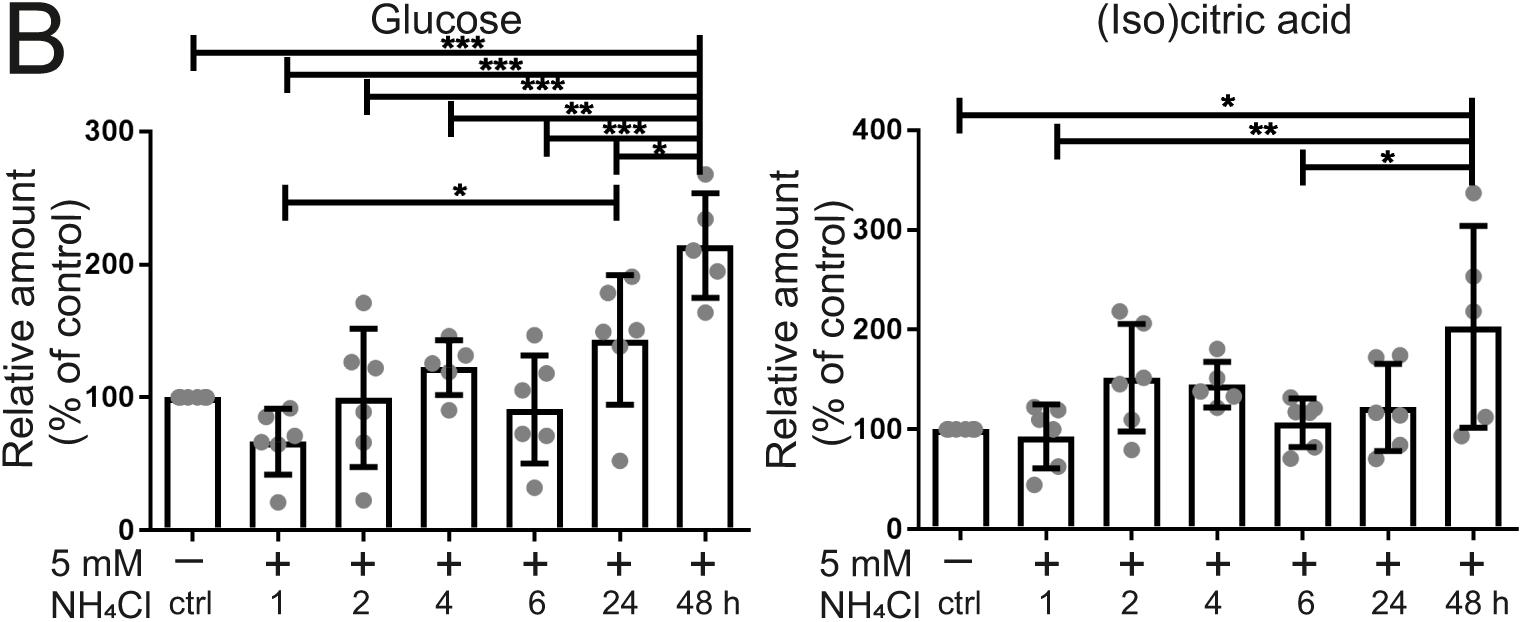
Ammonia alters energy and amino acid metabolism and anaplerotic precursor accumulation correlates with recovery. Metabolite abundances measured via GC-MS from human astrocytoma cells treated with 5 mM NH_4_Cl. Shown are relative abundances of amino acids (A) and glucose and (iso)citric acid (B) normalized to control (100%). Data represented as mean ± SD (n=4-6). Statistics: One-way ANOVA with Tukey’s post hoc test. *P < 0.05, **P < 0.01, ***P < 0.001.

### Targeted metabolomics analysis suggests alteration of the TCA-cycle induced by hyperammonemia

Ammonia leads to a rapid and drastic change in energy metabolism. To check whether metabolites linked to energy metabolism including amino acids and TCA-cycle intermediates are altered by ammonia treatment, we treated human astrocytoma cells with 5 mM NH_4_Cl for 1 to 48 h and subjected them to a targeted metabolomics analysis (Fig. 4AB and Fig. S4). We observed a significant increase in the level of isocitric and/or citric acid, which are indistinguishable by the GC-MS system used here, after 48 h of ammonia treatment. Moreover, branched chain amino acids (BCAAs), such as isoleucine/leucine/valine, are significantly increased after 24 h. Other essential amino acids methionine, phenylalanine, threonine and tryptophan were increased after 24 h and/or 48 h, respectively. The non-essential amino acids alanine, aspartate, cysteine, proline, serine and tyrosine were increased after 24 h and/or 48 h while glycine has a significant increase only after 4 h of treatment. We noted, that with the increase in numerous amino acids at late time points also a slight improvement of mitochondrial respiration is seen (Fig. 2B and Fig. S1A) which could point to a delayed role of these amino acids in a compensatory response. To test the fate of ammonia further, we traced incorporation of NH_4_^+^ in human astrocytoma cells using isotopically labeled ^15^NH_4_Cl under the same conditions as before. Using LC-MS analysis we found the strongest enrichment of ^15^N-isotopologues for glutamate, followed by aspartate, proline, and BCAAs (Fig. 5A-E and Fig. S5). Of note, a strong enrichment occurred already after 1 h, emphasizing a rapid effect of hyperammonemia. The enrichment of ^15^N-isotopologues was highest after 4 and 24 h. Interestingly, the synthesis of glutamate and its downstream metabolites aspartate and proline, as well as the BCAAs, are catalyzed by the enzyme glutamate dehydrogenase, followed by downstream reactions, *e.g.* transaminase reactions to form BCAAs (Fig. 5F). Overall, this pattern indicates that mitochondrial glutamate dehydrogenase 2 (GDH2, encoded by the *GLUD2* gene), which catalyzes the reductive amination of α-ketoglutarate and ammonia to glutamate, could be involved in the initial fixation of ammonia in mitochondria.

**Figure 5:**
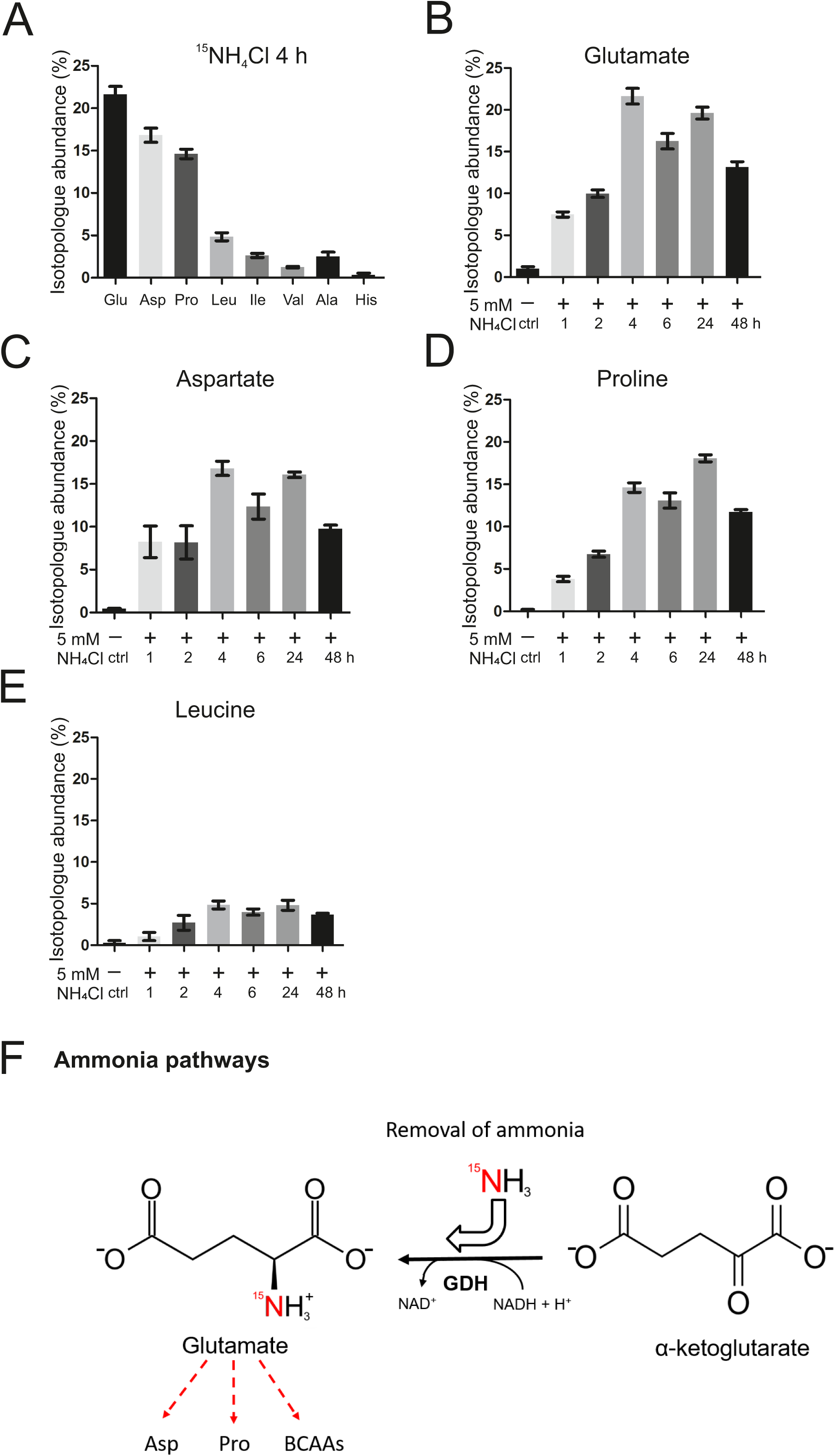
Ammonia is incorporated into GDH-dependent metabolites. Relative enrichment of ^15^N-labeled amino acids measured via LC-MS from human astrocytoma cells treated with ^15^NH_4_Cl. (A) Amino acids found to be enriched in ^15^N-isotopologue abundance, representative after 4 h treatment. Isotopologue abundance of ^15^N in glutamate (B), aspartate (C), proline (D) and leucine (E) over time. (F) Pathway of ^15^N-ammonia recycling and utilization via glutamate dehydrogenase (GDH) and secondary reactions. Red arrows indicate the metabolic path for nitrogen. Data represented as mean ± SD (n=3).

### Ammonia-induced impairment of mitochondrial respiration depends on GDH2

To test this hypothesis GDH2 was downregulated in human astrocytoma cells by targeting the *GLUD2* gene using siRNA. Knock-downs were validated using Western blot analysis (Fig. 6A, SFig. S6) and qPCR (SFig. S6). Cells depleted for GDH2 were subjected to analysis of OCR using the Mito Stress Test. Depletion of GDH2 efficiently prevented the reduction of mitochondrial respiration upon treatment for 1 h with ammonia (Fig. 6B). To investigate this further GDH2 was overexpressed (Fig. 6A) and subjected to OCR measurements. Treatment of GDH2-overexpressing cells with 5 mM NH_4_Cl for 1 h exacerbates the decrease of mitochondrial respiration observed before, with and without additional transfection with siGLUD2 (siRNA not targeting *GLUD2* plasmid sequence) (Fig. 6CD). In accordance with the GDH2 knock-down results presented before, the transfection with empty vector (ev) and siGLUD2 prevented NH_4_Cl-induced impairment of respiration, confirming our knock-down results described above (Fig. 6D). Taken together, we provide strong evidence that human glutamate dehydrogenase 2 is a critical factor for the rapid impairment of mitochondrial respiration caused by hyperammonemia.

**Figure 6:**
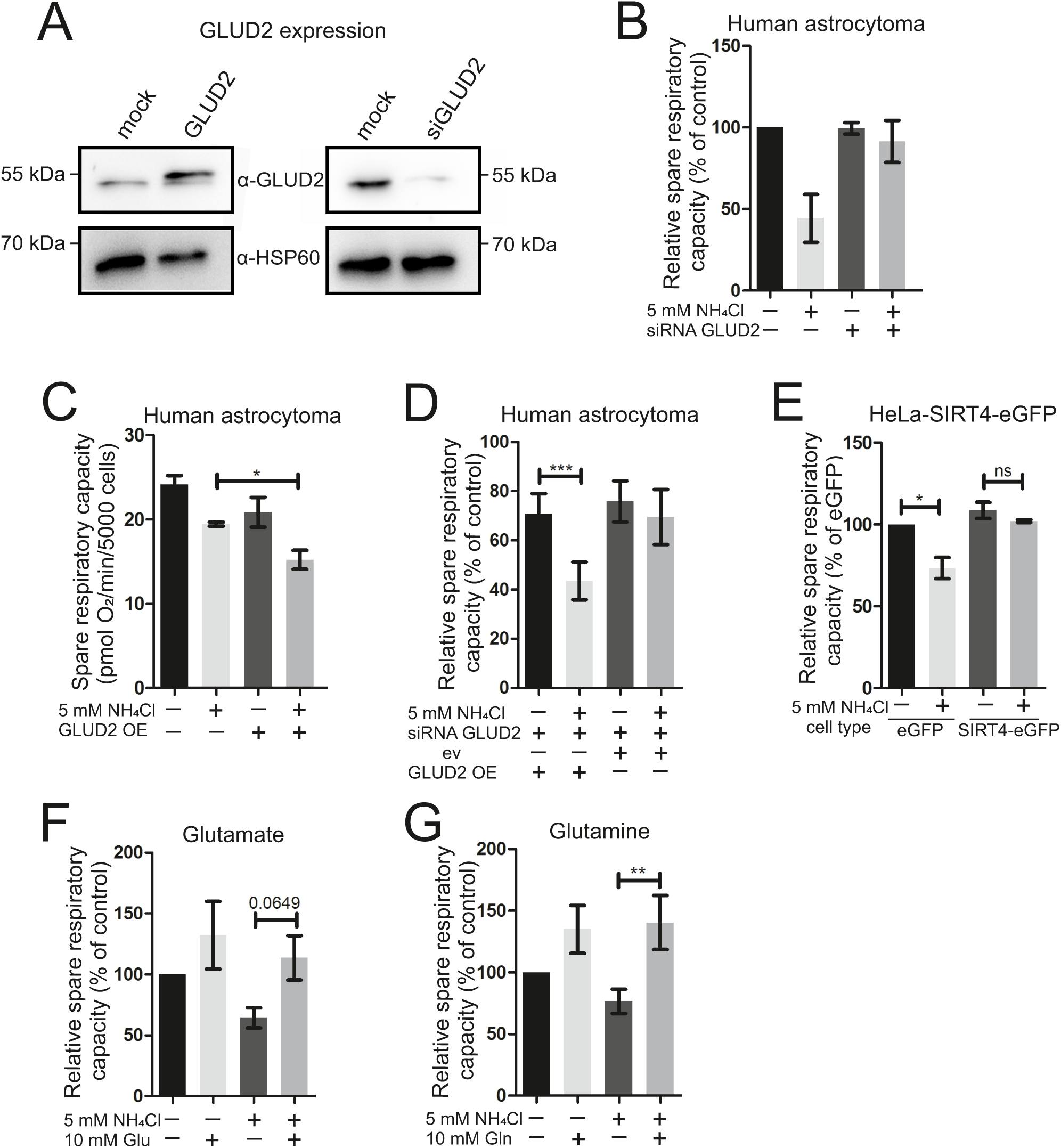
Knock-down of *GLUD2* and anaplerotic supplementation rescues, whereas overexpression of *GLUD2* exacerbates detrimental effect of NH_4_Cl on mitochondrial respiration. (A) Representative Western blot analysis of human astrocytoma cell lysates upon *GLUD2* overexpression (48 h) and knock-down (48 h) vs. mock samples. HSP60 was used as loading control. (B) Oxygen consumption rate (OCR) in human astrocytoma cells was analyzed on Seahorse XFe96 Extracellular Flux Analyzer with the Mito Stress Test Kit. After *GLUD2* knock-down cells were treated with 5 mM NH_4_Cl for 1 h. Individual biological replicates normalized to controls (100%). (C) Spare respiratory capacity of human astrocytoma cells. After *GLUD2* overexpression cells were treated with 5 mM NH_4_Cl for 1 h (n=5). (D) Relative spare respiratory capacity of human astrocytoma cells. Treatment was done with 5 mM NH_4_Cl for 1 h. GLUD2 expression from plasmid (GLUD2 OE) is not sensitive to the GLUD2 siRNA used. Individual biological replicates normalized to controls (100 %, not shown) (n=5). (E) Relative spare respiratory capacity measured with Seahorse Analyzer using Mito Stress Test Kit. HeLa-eGFP and HeLa-SIRT4-eGFP cells treated with 5 mM NH_4_Cl for 1 h were analyzed with respective controls. Individual biological replicates were normalized to HeLa-eGFP cells (100 %) (n=3). (F), (G) Human astrocytoma cells were subjected to Mito Stress Kit measurement on Seahorse XFe96 Extracellular Flux Analyzer. Cells were treated as indicated with compounds (10 mM) and 5 mM NH_4_Cl, or not, for 1 h. Relative spare respiratory capacity represented by oxygen consumption rate (OCR) was measured with glutamate (F) (n=4-5), glutamine or glutamine (G) (n=5). Individual biological replicates were normalized to controls (100%). Data are presented as mean ± SEM. Statistics: One-tailed student’s t-test comparing 2 groups. *P < 0.05, **P < 0.01, ***P < 0.001, ns: not significant.

**Figure 7:**
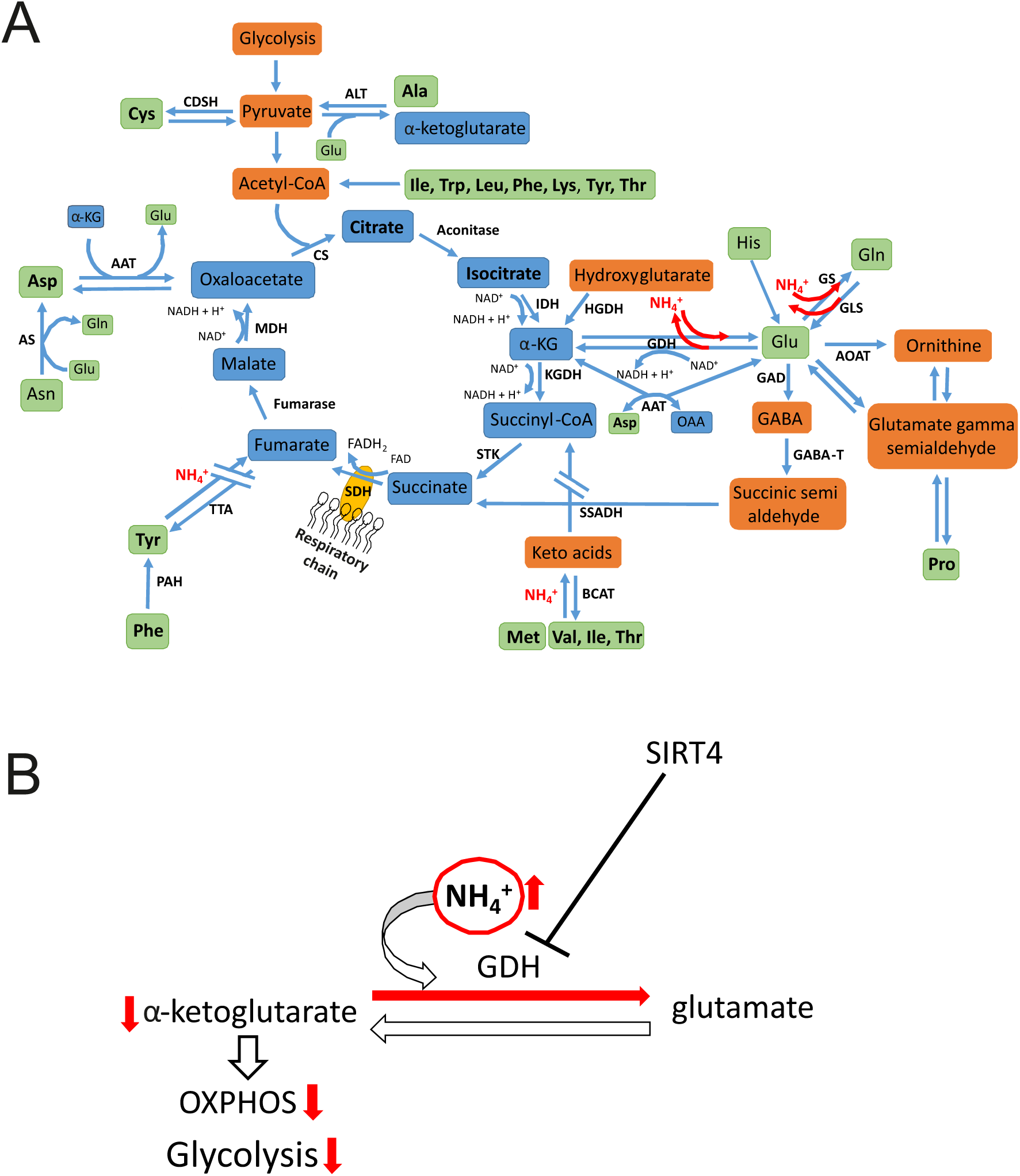
Proposed mechanism of influence of ammonia on mitochondrial metabolism. (A) Impact of excess of ammonia on the TCA-cycle and anaplerotic reactions. Amino acids and TCA-cycle intermediates increased in steady-state metabolites are depicted in bold. (B) Proposed mechanism on the role of glutamate dehydrogenase (GDH) on ammonia-induced toxicity. α-KG: α-ketoglutarate, AAT: aspartate aminotransferase, ALT: alanine aminotransferase, AOAT: acetyl ornithine aminotransferase, CDSH: Cysteine desulfhydrase, CS: citrate synthase, GABA-T: GABA-transaminase, GAD: glutamate decarboxylase, GDH: glutamate dehydrogenase, GLS: glutaminase, GS: glutamine synthetase, HGDH: hydroxyglutarate dehydrogenase, IDH: isocitrate dehydrogenase, KGDH: α-ketoglutarate dehydrogenase, MDH: malate dehydrogenase, OAA: oxaloacetate, PAH: phenylalanine hydroxylase, SDH: succinate dehydrogenase, SSADH: succinate semialdehyde dehydrogenase, STK: succinate thiokinase, TTA: tyrosine transaminase.

### Inhibition of GDH-dependent reductive amination of α-ketoglutarate by supplementation with glutamate or glutamine or by SIRT4 overexpression improves mitochondrial respiration under hyperammonemia

To further test the role of GDH in ammonia-induced toxicity on mitochondrial respiration, we aimed to rescue this detrimental effect using different amino acids. Pretreatment with 10 mM glutamine or glutamate directly before treatment with 5 mM NH_4_Cl for 1 h both led to an apparent rescue of ammonia-mediated decrease in mitochondrial respiration (Fig. 6FG). This would be consistent with the rapid ability to produce α-ketoglutarate directly from glutamate (via GDH or AATs) or indirectly by producing glutamate from excess of glutamine by mitochondrial glutaminase. GDH is further known to be inhibited by ADP-ribosylation catalyzed by the mitochondrial sirtuin SIRT4 (Haigis et al., 2006). We employed HeLa-SIRT4-eGFP cells, a stable cell line overexpressing SIRT4 and the corresponding control cells (HeLa-eGFP) to test whether inhibition of GDH2 by SIRT4 can also restore mitochondrial respiration in the presence of hyperammonemia. OCR of both cell types was measured with or without treatment with 5 mM NH_4_Cl for 1 h. The spare respiratory capacity of untreated HeLa-SIRT4-eGFP and HeLa-eGFP were not affected whereas after ammonia treatment respiration dropped significantly in HeLa-eGFP cells but not in cells overexpressing SIRT4 (Fig. 6E). Overall, several lines of evidence assign an essential role to the mitochondrial glutamate dehydrogenase, GDH2, for the rapid impairment of mitochondrial respiration under hyperammonemia. Mechanistically, we propose that GDH2 is required for the ammonia-induced depletion of α-ketoglutarate by reductive amination to glutamate (Fig. 7AB).

## Discussion

Hepatic encephalopathy is a common complication occurring upon severe liver dysfunction. Albeit it is largely accepted that hyperammonemia-induced impairment of astrocyte function plays a major role in mediating neurological disturbances (Görg et al., 2018; Häussinger and Görg, 2010; Norenberg, 1987), the underlying molecular mechanisms are not well understood and numerous models were proposed (Cash et al., 2010; Cordoba, 2014; Ott and Vilstrup, 2014). One particular aspect that is quite controversially discussed is the role of altered brain energy metabolism caused by hyperammonemia. Alterations of the metabolome of CSF of HE patients (Weiss et al.), differences in respiratory chain enzyme activities in rat HE models (Boer et al., 2009; Dhanda et al., 2017), and variable results on altered levels/activities of TCA-cycle metabolites and enzymes have been reported (Rama Rao and Norenberg, 2012). Overall, it is clear that the role of alteration of brain energy metabolism and alterations in the TCA-cycle is postulated, yet also contradicting views have been put forward. So far, it is not possible to explain these discrepancies. However, it could well be that different models and/or different time points of analysis can at least partly explain these results. We noticed that the large majority of studies investigating acute as well as chronic *in vitro* and *in vivo* HE models address toxic effects of hyperammonemia after one day, few days, or even after weeks (Görg et al., 2015; Hazell and Norenberg, 1998; Oenarto et al., 2016; Qvartskhava et al., 2015). Very little is known about the immediate effect of ammonia on brain energy metabolism. Here we provide several lines of evidence that high ammonia levels lead to a rapid major metabolic reprogramming of astrocytes, in particular via GDH-dependent impairment of the TCA-cycle, and we suggest that this represents an early event in the pathogenesis of HE.

The following arguments strongly support this view. (1) Our data show that ammonia introduces rapid mitochondrial fragmentation in two cell types, namely in human astrocytoma cells and primary rat astrocytes. Alterations of mitochondrial morphology are well known to occur upon mitochondrial dysfunction and upon metabolic reprogramming (Chan, 2006; Duvezin-Caubet et al., 2006; Zuchner et al., 2004). (2) In primary cells the effect was more rapidly observed than in human astrocytoma which could be attributed to metabolic differences known to exist in tumor cells (Hsu and Sabatini, 2008). (3) Hyperammonemia resulted in a virtually instantaneous impairment of mitochondrial respiration in primary rat astrocytes and human astrocytoma. (4) The inhibitory effect of ammonia on respiration was concentration dependent. (5) The effect was not mediated by possible changes in pH as a pH-mimetic compound did not impair respiration, excluding the immediate effect on mitochondrial function representing a secondary pH-mediated effect. (6) Glycolysis was likewise markedly hampered during hyperammonemia as a consequence of TCA-cycle inhibition. (7) Metabolomic analyses revealed that ammonia treatment resulted in an increase of several amino acids including those that are directly or indirectly able to engage in anaplerotic reactions feeding the TCA-cycle. (8) Glucose levels were increased consistent with the observation that glycolysis is inhibited rapidly as well by ammonia. (9) Isotope-labelling of ammonia and metabolic tracing showed that ammonia is rapidly fixed in glutamate (and metabolites derived from glutamate). This prompted us to investigate the role of the glutamate dehydrogenase which catalyzes the reductive amination of α-ketoglutarate and ammonia to glutamate. (10) Indeed, ammonia-induced inhibition of mitochondrial respiration is strongly dependent on the mitochondrial GDH2 suggesting that removal of ammonia may occur via GDH2. (11) This is further supported by the observation that GDH2 inhibition by SIRT4 overexpression has the same effect as GDH2 knock-down. (12) Moreover, GDH2 overexpression resulted in a sensitization of astrocytes to ammonia. (13) Glutamate as well as glutamine addition was beneficial suggesting that promoting anaplerosis of the TCA-cycle via GDH2 can reduce ammonia-induced toxicity. Taken together, we have several consistent lines of evidence that a critical entry point of excessive ammonia is the reductive amination of α-ketoglutarate by GDH2 causing impairment of the TCA-cycle and thus consequently of respiration as well as glycolysis. GDH2 is known to play an important part specifically in astrocytes, yet its role under hyperammonemia is not well understood. It was reported that *hGLUD2* expression in astrocytes increases the capacity for uptake and oxidative metabolism of glutamate, particularly during increased workload and aglycemia, implying GDH2 as an important mediator allowing to replenish the TCA-cycle from glutamate (Nissen et al., 2018). This is in line with our data showing that glutamate can indeed ameliorate the effect of hyperammonemia. A recent study showed that metabolic differences between transgenic *hGLUD2* mice and control mice during postnatal brain development center on metabolic pathways surrounding the TCA-cycle (Li et al., 2016). These results put another line of support for the importance of the GDH2-dependent modulation of energy metabolism in the brain.

In contrast to the prevailing view, our data show that under hyperammonemia α-ketoglutarate is primarily converted to glutamate and not vice versa. To our knowledge, this direction of the GDH reaction was not considered to be relevant for the pathogenesis of HE so far. This does not dispute the role of glutamate dehydrogenase as an important enzyme for anaplerosis, but GDH was not considered to be able to remove ammonia in the brain when concentrations are high enough to drive the reaction into the opposite direction. Still, an ammonia-detoxifying role of GDH was already shown to occur in the liver (Williamson et al., 1967) and was proposed based on a mathematical model employed to investigate the mechanism of ammonia detoxification in the liver (Ghallab et al., 2016). Consistent with this, injection of GDH and α-ketoglutarate with cofactors into mice led to a reduction of ammonia blood levels to normal levels within 15 minutes (Ghallab et al., 2016). Additionally, a rapid decrease in α-ketoglutarate concentration in rat liver was reported after injection of ammonium chloride solution (Williamson et al., 1967). To our knowledge, the role of GDH-dependent ammonia detoxification in other tissues than the liver was not addressed so far. Thus, our data now show that GDH2 in human astrocytes on one hand helps to remove the ammonia load, but on the other hand impairs the TCA-cycle. It acts as a double-edged sword as GDH2-mediated removal of ammonia is not only beneficial, as commonly expected, but rather detrimental to astrocytes. As a result of impairing the TCA-cycle we can well explain why glycolysis as well as OXPHOS is rapidly inhibited as a consequence of hyperammonemia. We see a clear concentration dependency of ammonia-induced effects on respiration. This is in accordance to most patient data showing a correlation between blood ammonia levels and severity of disease (Ong et al., 2003). Nevertheless, there are exceptions where ammonia levels are not directly correlated to the patient’s state of health, but reasons for this are still unknown (Kramer et al., 2000; Ong et al., 2003; Stahl, 1963).

It is interesting to note that humans and apes, in contrast to other mammals, have two genes encoding GDH, namely GDH1 and GDH2, encoded by GLUD1 and *GLUD2*, respectively. GDH1 is found in all mammals in the cytosol and mitochondria and is widely expressed in all tissues. On the other hand, *GLUD2* encodes a mitochondrial form of GDH that is expressed solely in testis, epithelial kidney cells and astrocytes of higher apes and humans. It is proposed that *GLUD2* appeared in evolution after retroposition of the GLUD1 gene, probably in an ape ancestor less than 23 million years ago (Burki and Kaessmann, 2004). The mature forms of human GDH1 (hGDH1) and hGDH2 are highly homologous with a sequence similarity of about 97% (Mastorodemos et al., 2009). Under physiological conditions, the GDH reaction mainly catalyzes the oxidative deamination to form α-ketoglutarate (Adeva et al., 2012). GDH is regulated by SIRT4, a mitochondrial enzyme that uses NAD to ADP-ribosylate and by that to inhibit GDH activity (Haigis et al., 2006). Here, we show that overexpression of SIRT4 can phenocopy the effect of GDH2 depletion by siRNA corroborating our results.

One argument allowing us to conclude that ammonia is utilized in the GDH reaction towards glutamate was obtained from ^15^N-label incorporation mainly in Glu, Asp, and Pro as well as to a lower extent in Leu, Ile, Val, Ala, and His after administration of ^15^NH_4_Cl. Glu, Pro, Asp and the BCAAs are directly associated with GDH downstream reactions or acquire the ^15^N-label through secondary reactions. It is interesting to note that also breast cancer cells were reported to fix ammonia via the GDH reaction (Spinelli et al., 2017). This study reported that in breast cancer cell lines ammonia is primarily assimilated through reductive amination catalyzed by GDH and the high accumulation of labeled nitrogen in Glu, Pro, Asp and Ala, among others, is strongly reminiscent to our data for astrocytes.

Other studies support the critical role of the TCA-cycle in HE. Weiss and colleagues published data examining metabolomics to highlight dysfunctions of metabolic pathways in CSF samples of HE patients (Weiss et al., 2016). This study revealed an accumulation of acetylated compounds, which also points towards a defect in the TCA-cycle. Additionally, the metabolites that were increased are involved in ammonia, amino acid and energy metabolism, for example glutamate, glutamine, methionine, phenylalanine and others (Weiss et al., 2016); many of which we also found elevated in our study. Patient data show an association between arterial hyperammonemia and increase of glutamine concentration in the brain (Tofteng et al., 2006), which could result from GDH activity towards glutamate and subsequently glutamine. It was further shown that under normal conditions GDH is important to sustain the catalytic activity of the TCA-cycle in mouse astrocytes by mediating the net formation of TCA-intermediates and that reduced GDH expression induces the usage of alternative substrates such as BCAAs (Nissen et al., 2015).

On the contrary, we show that when ammonia levels are high with increased time of treatment the concentration of amino acids such as BCAAs increases. How can that be explained or why is the effect of ammonia on mitochondrial respiration most prevalent at short time points but becomes less pronounced with time? We attribute this to a compensatory mechanism that involves the induction of anaplerotic reactions other than the one catalyzed by GDH. Such a mechanism could be enhanced proteolysis by autophagy or by proteasomal degradation of proteins. Indeed, autophagy was found to be induced in the *substantia nigra* of mice with liver damage and subsequent hyperammonemia (Bai et al., 2018) and in *Ulk1/2* DKO mouse embryonic fibroblasts (MEF) treated with 2 mM NH_4_Cl for 24 h (Cheong et al., 2011). In our study, when ammonia was washed-out for only one hour, the detrimental effect on mitochondrial oxygen consumption was not only gone, but respiration rather even reached levels above control levels, in particular after longer pretreatments with ammonia. This could be due to the fact that accumulated amino acids can rapidly engage in anaplerotic reactions and drive the TCA-cycle as soon as ammonia is removed. Thus, accumulated amino acids are “fired” into the TCA-cycle leading to an increase in respiration. Further, detrimental effects of ammonia are rapidly reversible which was also seen in mitochondrial morphology. HE has long been considered to be a reversible condition after liver transplantation, albeit a persistence in cognitive impairment in patients with preceding episodes of overt HE have been reported as well (Bajaj et al., 2010).

Hohnholt and colleagues could also show the importance of GDH for mitochondrial respiration. In Cns-*Glud1^-/-^* mice (GDH1 knock-out in synaptosomes) the basal respiration in brain mitochondria in presence of glutamate and malate was significantly reduced. In GDH1 knock-out neurons they show that without stimulation of respiration (e.g. by FCCP) there is no effect of GDH1 knock-out on respiration, while upon stimulation by FCCP the cells do not respond with an increased respiration (Hohnholt et al., 2017). This is in line with our results showing that hyperammonemia does not grossly affect basal mitochondrial respiration but rather when respiratory activity is induced *e.g.* by dissipation of the membrane potential. We suppose that this is an interesting aspect as it implies that astrocytes are more severely impaired under hyperammonemia when mitochondrial energy conversion is increased which could be the case under increased neuronal activity.

Our findings suggest a new mechanism in the pathogenesis of hepatic encephalopathy, involving all pathways of energy metabolism with a special emphasis on the role of GDH and its negative regulation by SIRT4. These results could open new strategies on the treatment of HE and some therapeutical instructions might have to be reconsidered. For example, for decades patients were advised to reduce dietary protein intake in order to avoid ammonia generation in the gastrointestinal tract through bacterial digestion (Cordoba, 2014). Fortunately, this practice has been revoked as weakening patients through protein restriction is not beneficial for recovery. Reducing the amount of amino acids could even increase the effects of ammonia-induced toxicity. New treatment options by supplementation of amino acids, e.g. glutamate, could help to reduce symptoms. Targeting sirtuins is already studied intensively regarding cancer research. However, this only holds true for SIRT1-3 and 5. For these sirtuins activators and inhibitors are known and studied extensively as potential cancer treatment options (Hu et al., 2014). SIRT4 is regulated by miR15a/b (Lang et al., 2016), yet a specific compound inhibiting SIRT4 activity is not known so far. Targeting SIRT4 could present a very promising future therapeutic target in novel, future treatments of hepatic encephalopathy. Further studies, especially on the potential role of SIRT4 and GDH regulation are needed for the development of improved treatment strategies of HE symptoms.

## Acknowledgements

We thank Prof. Dr. Dieter Häussinger and Dr. Boris Görg for providing primary rat astrocytes and continuous support, and Prof. Wilhelm Stahl and Prof. Peter Brenneisen for scientific discussions. We thank Andrea Borchardt and Tanja Portugall for excellent technical assistance. We are grateful to Dr. Helmut Hanenberg for providing cloning vectors. We appreciate the excellent technical assistance by Katrin Weber, Elisabeth Klemp, and Maria Graf in mass spectrometric analysis.

---

**Figure S1:**
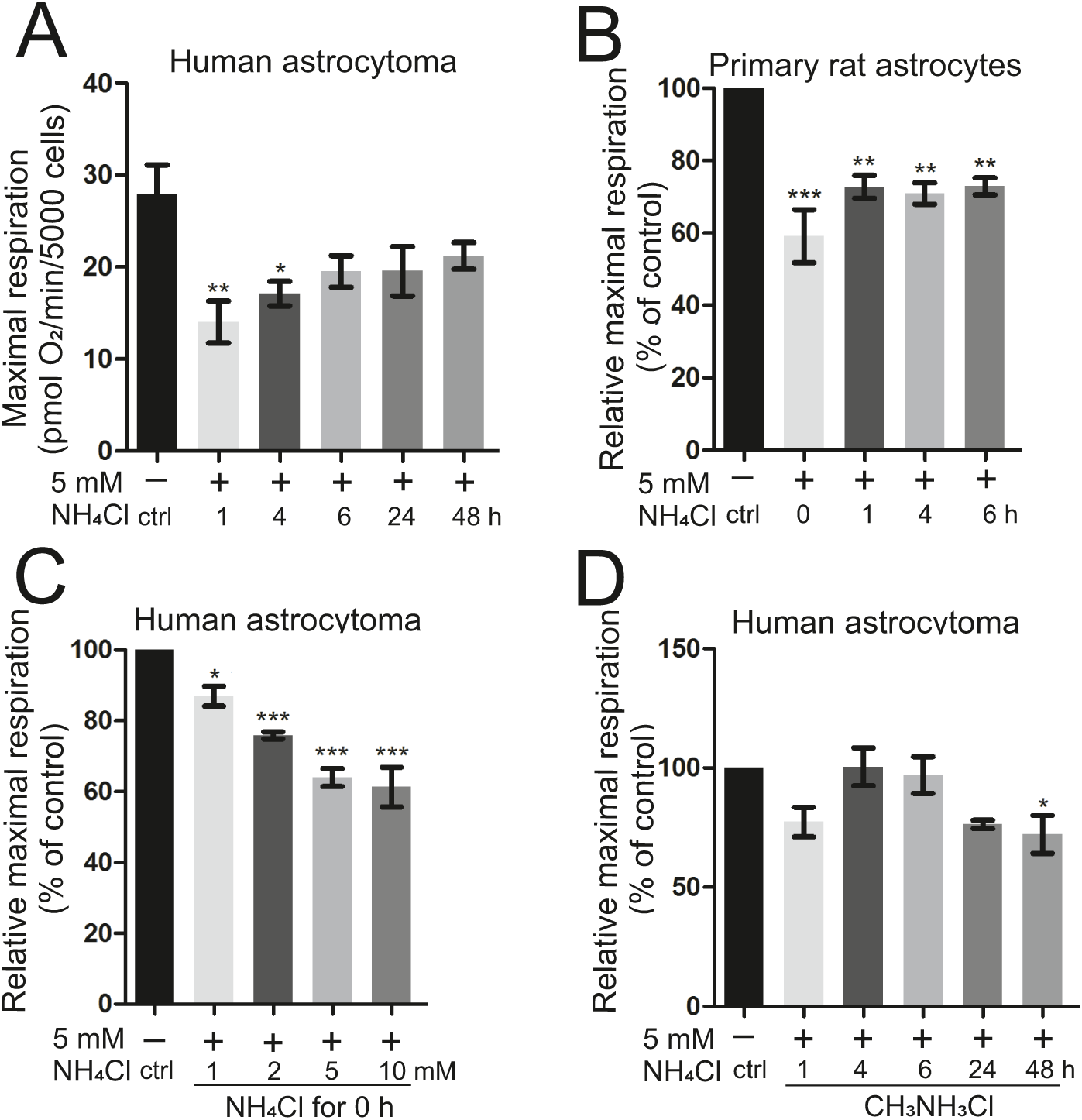
Mitochondrial respiration is immediately impaired by ammonia in a pH-independent manner. Oxygen consumption rate (OCR) of human astrocytoma cells and primary rat astrocytes was analyzed in Seahorse XFe96 Extracellular Flux Analyzer with the Mito Stress Test Kit after treatment with ammonia at indicated molarities or durations. (A) Maximal respiration of human astrocytoma cells after treatment with 5 mM NH_4_Cl for 1-48 h (n=5-7). (B) Relative maximal respiration of primary rat astrocytes after treatment with 5 mM NH_4_Cl for 1-6 h and directly after treatment (0 h) (n=3-4). (C) Relative maximal respiration of human astrocytoma cells and directly after treatment (0 h) with 1, 2, 5, or 10 mM NH_4_Cl (n=3). (D) Relative maximal respiration of human astrocytoma cells was determined after treatment with 5 mM CH_3_NH_3_Cl (pH-mimetic) for 1-48 h (n=3). (B), (C), (D) Individual biological replicates normalized to control (100%) are depicted. Data presented as mean ± SEM. Statistics: One-way ANOVA with Dunnett’s post test. *P < 0.05, **P < 0.01, ***P < 0.001.

**Figure S2:**
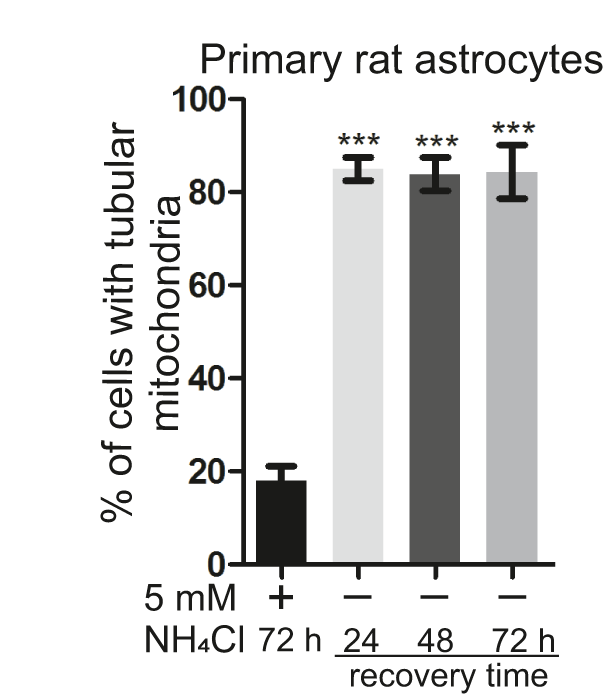
Ammonia-induced mitochondrial fragmentation is reversible. Primary rat astrocytes transfected with pEGFP-Mito. Baseline quantification was performed after 72 h treatment with 5 mM NH_4_Cl. Other time points represent recovery time periods after removal of ammonia. Characterization of mitochondria with respect to fragmented versus tubular morphology. Data represented mean ± SD (n=3). One-way ANOVA with Dunnett’s post test. ***P < 0.001.

**Figure S3:**
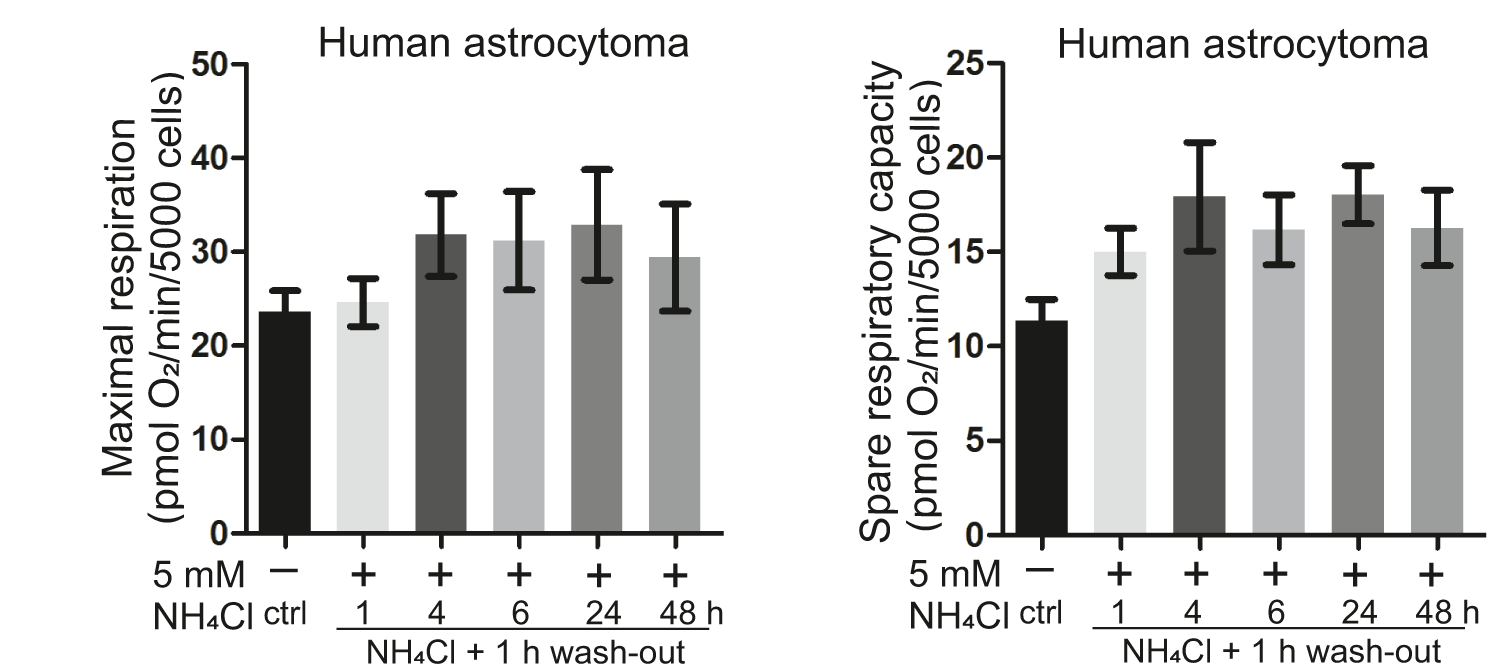
Ammonia-induced decrease in oxygen consumption rate is rapidly reversible. Human astrocytoma cells were analyzed using the Mito Stress Test Kit on Seahorse XFe96 Extracellular Flux Analyzer. Oxygen consumption rate (OCR) of maximum respiration (left) and spare respiratory capacity (right) were determined after 1 h wash-out of ammonia treated with 5 mM NH_4_Cl for 1 to 48 h. Data represent mean ± SEM (n=3).

**Figure S4:**
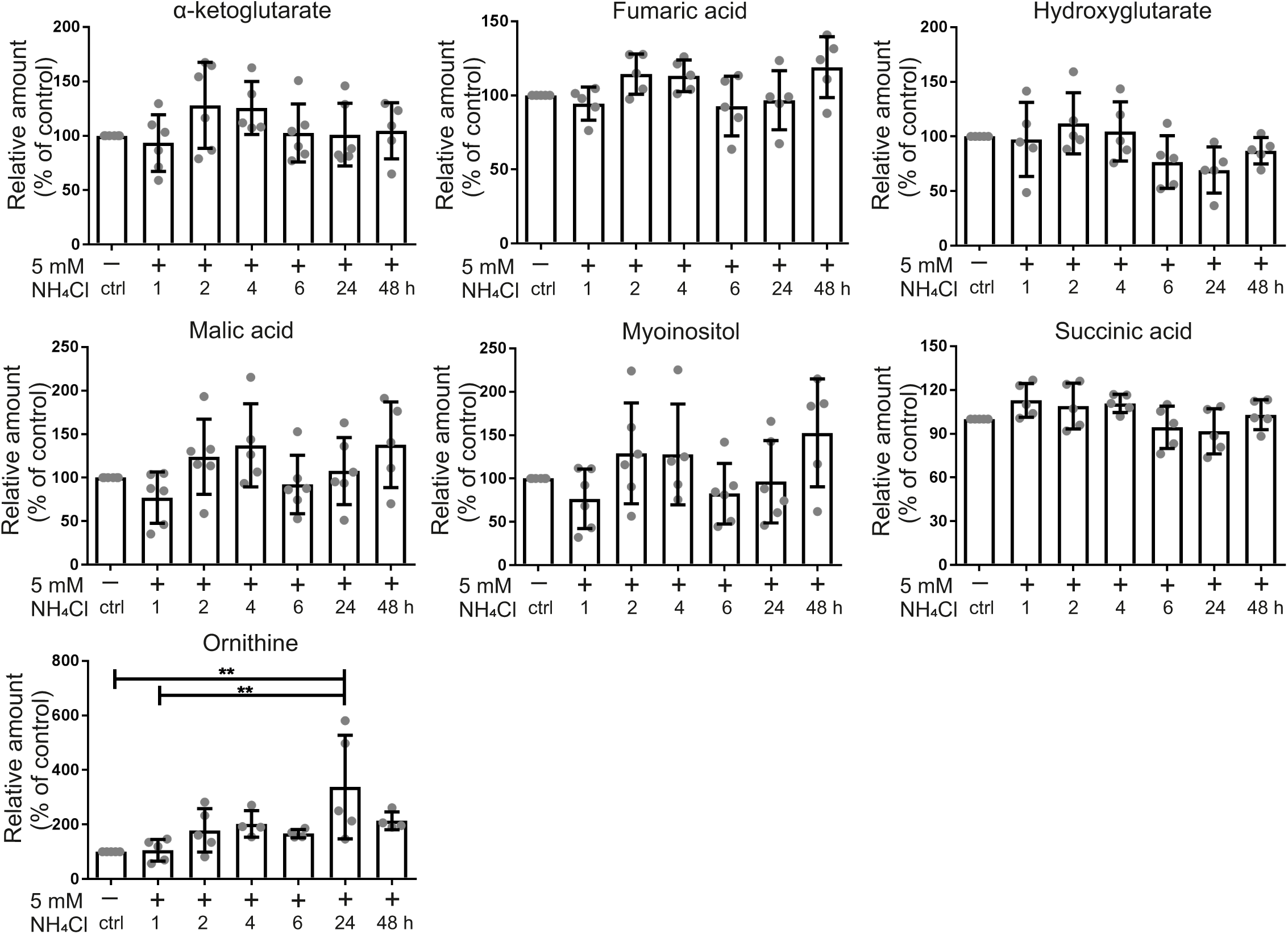
Additional results from steady-state metabolomics. Mass Spectrometry for steady-state metabolites was done in human astrocytoma cells on GC-QTOF. Treatment with 5 mM NH_4_Cl for 1-48 h. Relative abundance of respective metabolites compared to controls (100 %) over time. Data represent mean ± SD (n=4-6). Statistics: One-way ANOVA with Tukey’s post test. **P < 0.01.

**Figure S5:**
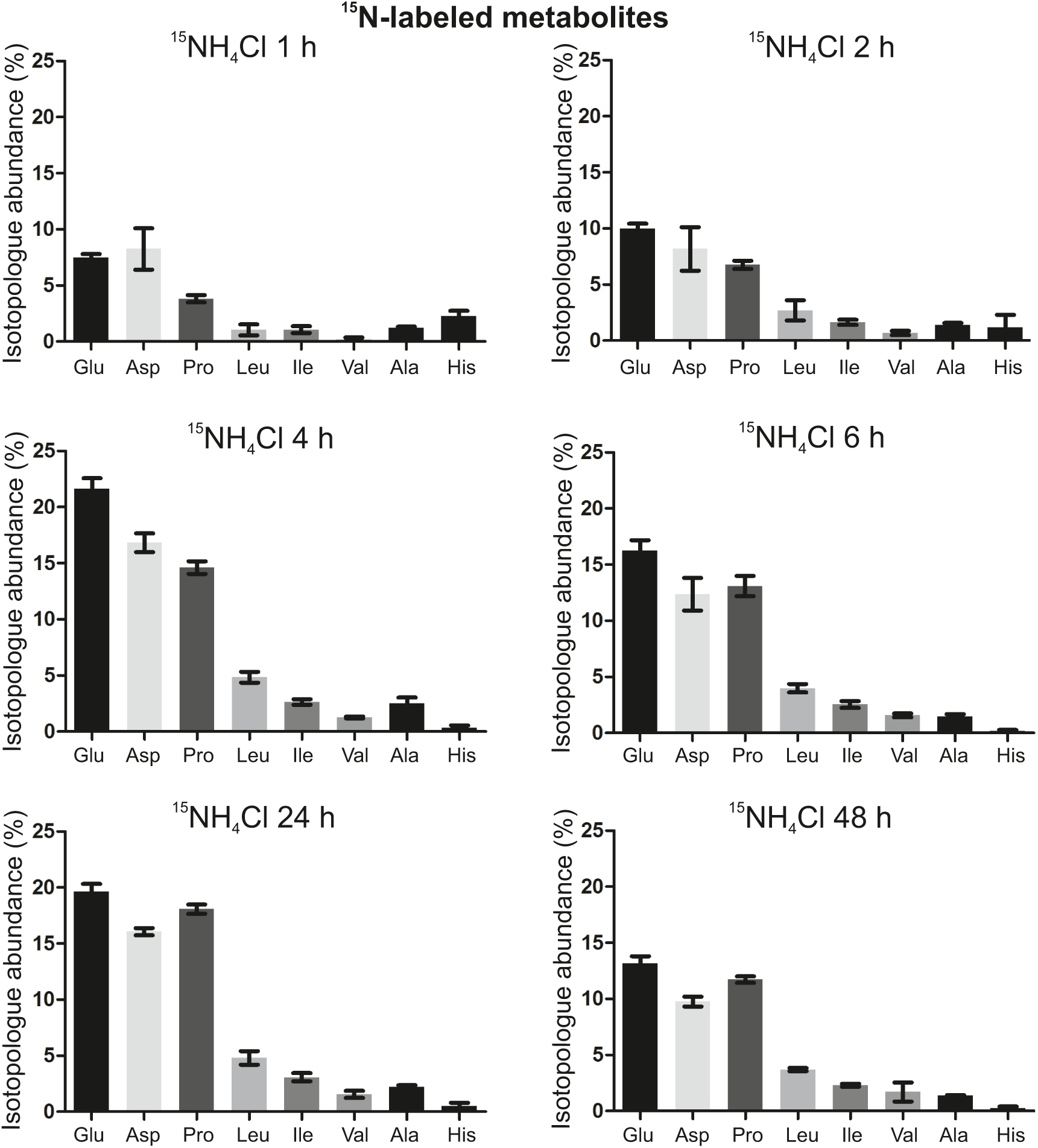
^15^N-isotopologue abundance of labeled amino acids over time. Mass Spectrometry for ammonia flux was done in human astrocytoma cells on LC-QTOF. Cells were treated with ^15^NH_4_Cl for 1-48 h and isotopologue abundance of ^15^N in Glu, Asp, Pro, Leu, Ile, Val, Ala and His was determined over time. Data represent mean ± SD (n=3).

**Figure S6:**
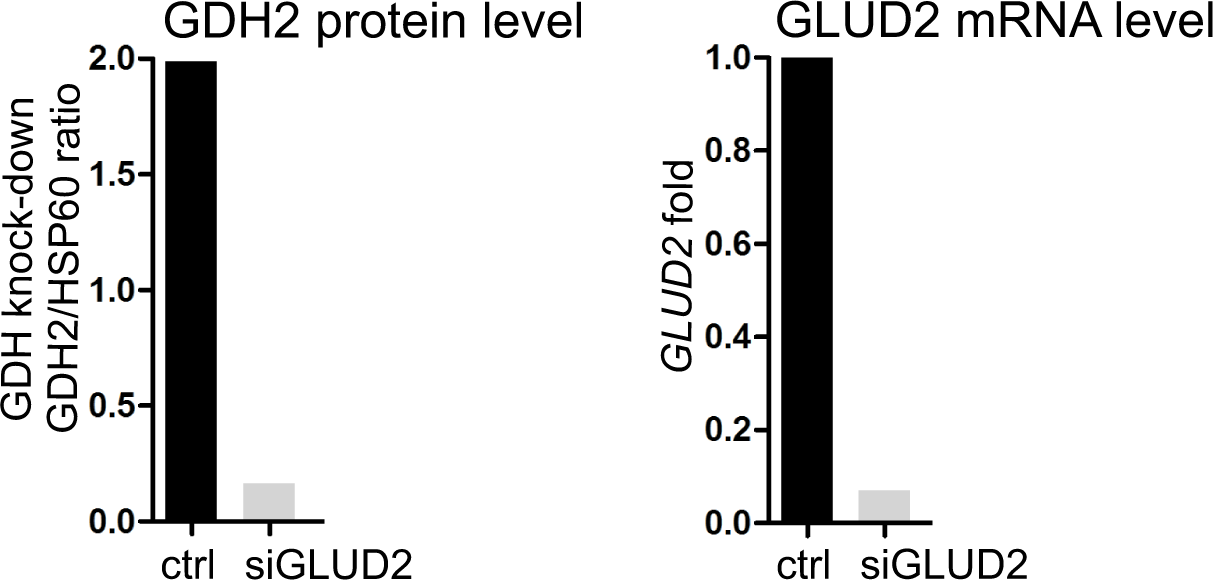
Protein and mRNA level of glutamate dehydrogenase after *GLUD2* knock-down. *GLUD2* knock-down validation in human astrocytoma cells. Left: Densitometry of protein levels determined by Western blot analysis (Fig. 6A) using HSP60 as loading control. Ratios of GDH2 to HSP60 levels are shown in knock-down *vs*. control cells. Right: mRNA level were determined by qPCR using HPRT1 as housekeeping gene. *GLUD2* expression levels are depicted in in knock-down cells as compared to control and normalized to 1. Representative experiments are shown.

