## Supplemental Tables 1 to 3 for "Ammonia inhibits energy metabolism in astrocytes in a rapid and GDH2-dependent manner"

**Supplemental Table S1. Raw data for targeted metabolite abundances measured via GC-MS from human astrocytoma cells treated with NH_4_Cl for 1 to 48 hours or not (control).**

|  | **Alanine** | **Valine** | **Leucine** | **Isoleucine** | **Proline** | **Glycine** | **Succinic acid** | **Fumaric acid** | **Serine** | **Threonine** | **Malic acid** | **Methionine** | **Aspartic acid** | **Cysteine** |
| --- | --- | --- | --- | --- | --- | --- | --- | --- | --- | --- | --- | --- | --- | --- |
| **control_1** | 123.0282066 | 18.9902447 | 15.0933263 | 10.2495073 | 1.08184889 | 62.7542946 |  |  | 41.8221387 | 47.3404029 | 21.6773031 | 5.7252943 | 108.2855737 | 1.35899103 |
| **control_2** | 82.97457134 | 16.3380233 | 21.0935423 | 14.0284994 | 5.6429083 | 9.44571045 | 4.946868274 | 10.14652495 | 27.2109555 | 33.4481314 | 5.97195766 | 6.32663334 | 59.74423847 | 2.57441821 |
| **control_3** | 142.9821127 | 26.204598 | 31.4617314 | 19.5112202 | 10.2114386 | 19.7211262 | 4.967354988 | 10.63390673 | 21.1035735 | 41.176682 | 8.39829423 | 6.16662224 | 85.86745617 | 0.75713076 |
| **control_4** | 10.63989651 | 1.92692013 | 2.35237755 | 1.54077365 | 1.26079184 | 0.64434518 | 1.059071043 | 1.532604537 | 1.82952716 | 2.46674063 | 0.84682394 | 0.43191033 | 7.243406381 | 0.20234494 |
| **control_5** |  |  |  |  |  |  |  |  |  |  |  |  |  |  |
| **control_6** |  |  |  |  |  |  |  |  |  |  |  |  |  |  |
| **1 h_1** | 50.41471022 | 15.1504547 | 10.5893385 | 8.55595203 | 0.95509926 | 27.5502227 |  |  | 23.9769184 | 27.1545639 | 7.66037023 | 2.04833012 | 16.32639389 | 0.17565713 |
| **1 h_2** | 82.80565467 | 22.4251086 | 26.5799984 | 19.6369346 | 3.7668592 | 13.6867071 | 6.274217869 | 10.27727373 | 70.7627802 | 60.9906 | 5.07559794 | 12.6157302 | 62.00194455 | 3.72578649 |
| **1 h_3** | 142.6080348 | 25.3253943 | 29.7782135 | 24.2909216 | 4.70256568 | 34.031474 | 6.11223995 | 9.805692686 | 67.8072899 | 53.4476391 | 7.35040833 | 3.66044644 | 31.43957265 | 0.37228066 |
| **1 h_4** | 7.096751736 | 1.05778624 | 1.27594187 | 0.99004823 | 0.73566035 | 1.01979001 | 1.066062106 | 1.501583292 | 1.94091311 | 2.22716493 | 0.88898944 | 0.43319368 | 9.355438751 | 0.23246101 |
| **1 h_5** | 2.884043763 | 0.6748289 | 0.79812431 | 0.55206083 | 0.21402619 | 0.43450215 | 1.101227464 | 1.169275362 | 0.9065547 | 0.98471558 | 0.39055053 | 0.19259829 | 3.531278416 | 0.15012796 |
| **1 h_6** | 7.720849048 | 1.00331368 | 1.09097806 | 0.81744562 | 0.40183979 | 1.0060165 | 1.166208172 | 1.604612507 | 2.1066592 | 2.57563644 | 0.87686022 | 0.48464084 | 8.644684156 | 0.22915123 |
| **2 h_1** | 102.2033027 | 28.1486347 | 22.9729096 | 18.5852905 | 3.03217716 | 58.130615 |  |  | 52.1751769 | 57.4233111 | 12.773316 | 5.01964414 | 48.39038899 | 0.62612042 |
| **2 h_2** | 124.886084 | 41.1670308 | 55.1611584 | 43.370623 | 10.8975833 | 16.9657537 | 6.128745864 | 12.95877916 | 118.513618 | 102.815492 | 7.92185073 | 21.4246429 | 95.1226066 | 8.19937328 |
| **2 h_3** | 153.9321692 | 31.0977781 | 38.0112758 | 31.365995 | 7.23832531 | 38.8943916 | 6.264234483 | 12.21646666 | 80.2271309 | 50.5219794 | 9.68986778 | 1.9468305 | 15.51322358 | 0.35619292 |
| **2 h_4** | 13.9997385 | 5.60991641 | 9.08138997 | 5.53927888 | 4.81913274 | 2.06120646 | 1.135434969 | 1.954174634 | 3.47331201 | 4.86207425 | 1.63634655 | 1.06167393 | 15.75363136 | 0.51792383 |
| **2 h_5** | 7.254819969 | 2.17171849 | 3.22779827 | 2.0279897 | 1.57762981 | 1.06704081 | 0.973862016 | 1.494723814 | 2.31110854 | 2.72889635 | 0.96305599 | 0.61045818 | 10.42797967 | 0.24668971 |
| **2 h_6** | 9.477525142 | 2.09557084 | 2.33879198 | 1.54447573 | 1.37193488 | 1.46665943 | 1.01576461 | 1.598096435 | 2.31241225 | 3.09682554 | 1.10356759 | 0.73609736 | 10.9699974 | 0.30811487 |
| **4 h_1** |  |  |  |  |  |  |  |  |  |  |  |  |  |  |
| **4 h_2** | 131.7298346 | 39.0900594 | 50.0312918 | 40.3029718 | 8.96033444 | 22.9702324 | 5.783683795 | 12.3059182 | 115.160659 | 97.2898668 | 7.56226523 | 21.2978601 | 74.7906489 | 5.45760186 |
| **4 h_3** | 153.2482137 | 37.0627015 | 50.041558 | 40.4163342 | 10.4612243 | 43.0603352 | 5.792090758 | 10.76024473 | 99.2996114 | 77.300178 | 8.95060695 | 9.36787183 | 46.78468022 | 1.162878 |
| **4 h_4** | 12.75942363 | 1.85045297 | 2.48201522 | 1.94838652 | 1.04903041 | 1.80187248 | 1.080037526 | 1.742552087 | 4.09934522 | 4.1147104 | 1.21906357 | 0.72070176 | 11.58393529 | 0.32535383 |
| **4 h_5** | 21.50495671 | 4.00018709 | 5.07727575 | 3.76818554 | 2.79566041 | 3.01620203 | 1.173298665 | 1.932389193 | 6.40782343 | 6.89689705 | 1.82271958 | 1.36255168 | 22.01388731 | 0.52178493 |
| **4 h_6** | 7.036885051 | 1.43550477 | 1.89467571 | 1.43422329 | 0.79988668 | 1.17236718 | 1.143746075 | 1.595734522 | 2.0649623 | 2.69073133 | 0.79193632 | 0.5684671 | 8.736893836 | 0.47599104 |
| **6 h_1** | 169.1879083 | 61.4450126 | 63.5134412 | 41.8731061 | 5.71161463 | 90.6499145 |  |  | 110.080182 | 115.373835 | 18.679939 | 7.07959984 | 71.43039313 | 0.98867755 |
| **6 h_2** | 149.7540449 | 44.7087035 | 59.7099686 | 48.9924978 | 10.6929225 | 19.3325755 | 5.446566015 | 11.132173 | 142.425163 | 135.556929 | 5.91285607 | 27.120992 | 89.75918548 | 6.40006075 |
| **6 h_3** | 92.02859884 | 71.692002 | 102.431268 | 62.4498718 | 23.8088954 | 34.5865882 | 5.239460342 | 9.04634661 | 51.5981328 | 59.2866037 | 6.2627912 | 10.9140218 | 48.33881157 | 0.92399044 |
| **6 h_4** | 8.49977045 | 1.60893414 | 1.9874244 | 1.49158646 | 0.67217035 | 1.03507186 | 1.02953361 | 1.412792778 | 2.15357338 | 2.78417181 | 0.74158267 | 0.64195118 | 9.201124336 | 0.26582709 |
| **6 h_5** | 18.06728798 | 2.47029302 | 3.06757993 | 2.33127959 | 1.40976475 | 1.41330996 | 0.880683865 | 1.737606016 | 6.09251405 | 5.86106317 | 1.29432807 | 1.1799676 | 15.09710321 | 0.41040259 |
| **6 h_6** | 4.316561962 | 1.51850072 | 2.2902991 | 1.47315348 | 1.02296762 | 0.50289848 | 0.80638215 | 0.976933237 | 1.74284042 | 1.69300741 | 0.4477923 | 0.39015684 | 4.795330929 | 0.17194198 |
| **24 h_1** | 253.517088 | 67.6734107 | 59.1776964 | 42.2122979 | 9.78541543 | 108.036883 |  |  | 119.699841 | 119.096295 | 20.6551496 | 9.09871896 | 147.6237107 | 2.70102798 |
| **24 h_2** | 124.4779877 | 39.0383471 | 49.8823855 | 40.353536 | 10.6530934 | 14.9799416 | 5.278848913 | 10.05291188 | 79.9049342 | 73.7965243 | 6.34893999 | 14.3569567 | 78.83140209 | 4.9519445 |
| **24 h_3** | 159.4361512 | 38.7173541 | 50.3242129 | 41.6760546 | 12.5855306 | 39.5966279 | 5.400239681 | 10.04901805 | 81.5633705 | 71.784994 | 7.86274161 | 11.5604284 | 82.22867455 | 1.88752528 |
| **24 h_4** | 15.91113771 | 4.3195976 | 6.3952876 | 4.33043973 | 3.16641575 | 1.7830928 | 0.937180732 | 1.518184039 | 5.26077253 | 5.34696728 | 1.15329579 | 1.29478922 | 15.0271014 | 0.59987876 |
| **24 h_5** | 4.621677097 | 1.24173738 | 1.51234862 | 1.02950963 | 0.42144626 | 0.47583468 | 0.780863116 | 1.034423839 | 1.22999165 | 1.8337569 | 0.43247703 | 0.46791846 | 4.903690263 | 0.25183128 |
| **24 h_6** | 17.18433712 | 8.50049464 | 14.016842 | 8.66537193 | 7.71311624 | 1.97797473 | 0.855926546 | 1.893465165 | 7.72954628 | 7.59765382 | 1.38022006 | 1.63873292 | 19.51709194 | 0.69619847 |
| **48 h_1** |  |  |  |  |  |  |  |  |  |  |  |  |  |  |
| **48 h_2** | 101.0071797 | 23.6327051 | 35.2801889 | 26.2910483 | 10.9974003 | 10.9527478 | 5.558258005 | 8.925370191 | 52.4273399 | 49.7053468 | 4.18738321 | 8.42361178 | 66.99879012 | 3.99384329 |
| **48 h_3** | 218.4093879 | 35.0827189 | 46.5392264 | 35.5086016 | 16.9252344 | 52.0132906 | 5.629741155 | 11.79554252 | 83.4430582 | 79.8229523 | 9.30836675 | 12.311498 | 147.7369045 | 3.14339783 |
| **48 h_4** | 26.94782892 | 6.83873545 | 8.5924152 | 5.97546283 | 4.13324006 | 1.91501336 | 1.068428558 | 1.903990296 | 6.84919294 | 8.45231575 | 1.18960164 | 1.32644831 | 15.74686619 | 0.52740933 |
| **48 h_5** | 31.57451241 | 5.23564606 | 5.68301334 | 4.43415754 | 2.06627976 | 1.27543034 | 1.062422061 | 2.158332641 | 8.62777144 | 10.2971416 | 1.61813274 | 1.62982755 | 20.62026943 | 0.852086 |
| **48 h_6** | 31.01147827 | 4.73202699 | 5.60936608 | 4.4597073 | 2.32410154 | 1.07449776 | 0.935918945 | 2.016896194 | 9.67334412 | 10.7273017 | 1.49407544 | 1.68500976 | 18.01380974 | 0.67970942 |

|  | **Hydroxyglutarate** | **alpha-Ketoglutarate** | **Glutamic acid** | **Phenylalanine** | **Ornithine** | **(Iso)citric acid** | **Tyrosine** | **Glucose** | **Myoinositol** | **Tryptophan** | **Lysine** |
| --- | --- | --- | --- | --- | --- | --- | --- | --- | --- | --- | --- |
| **control_1** |  | 1.481532173 | 75.02991881 | 3.336466414 |  | 29.06574014 | 13.9541871 | 7.31380508 | 745.06328 | 1.56837445 |  |
| **control_2** | 0.663273418 | 0.929837135 | 105.0907506 | 8.155869565 | 0.14363838 | 31.48887544 | 5.94612621 | 0.66362598 | 275.033326 | 2.43032729 |  |
| **control_3** | 0.895203729 | 0.91331539 | 115.4567683 | 6.703557324 | 0.09815 | 28.45573696 | 9.68941622 | 0.63599908 | 479.69046 | 2.51937664 |  |
| **control_4** | 0.08869127 | 0.087670324 | 10.53002174 | 0.654453846 | 0.0113115 | 2.574017853 | 0.97526103 | 0.0588021 | 21.5358258 | 0.31426779 | 0.04128664 |
| **control_5** |  |  |  |  |  |  |  |  |  |  |  |
| **control_6** |  |  |  |  |  |  |  |  |  |  |  |
| **1 h_1** |  | 0.874525322 | 1.713468427 | 0.692670637 |  | 12.93920064 | 4.91017922 | 1.54293086 | 240.09568 | 0.88693801 |  |
| **1 h_2** | 0.939629981 | 1.205318004 | 118.5263226 | 14.26488793 | 0.21099612 | 38.58983418 | 11.770395 | 0.42831155 | 254.404863 | 5.06234702 |  |
| **1 h_3** | 0.998370959 | 0.943037928 | 25.17690187 | 2.274230937 | 0.05588459 | 33.97066901 | 12.4273106 | 0.45180717 | 327.831604 | 1.56983371 |  |
| **1 h_4** | 0.079400428 | 0.075940893 | 10.02511471 | 0.521945907 | 0.01526373 | 2.574956396 | 0.64005519 | 0.05401995 | 24.1073428 | 0.29202094 | 0.03630244 |
| **1 h_5** | 0.043231119 | 0.062665514 | 3.585106365 | 0.272550603 | 0.00797316 | 1.616924408 | 0.25387715 | 0.05021021 | 9.31510675 | 0.11903246 | 0.02464223 |
| **1 h_6** | 0.084232202 | 0.096578671 | 12.39669637 | 0.682853227 | 0.01352559 | 2.825480181 | 0.60787851 | 0.03900494 | 23.9975134 | 0.2047753 | 0.06627042 |
| **2 h_1** |  | 1.168924221 | 5.826970397 | 1.485865499 |  | 23.07839058 | 10.9959859 | 1.66195862 | 421.263776 | 1.36138324 |  |
| **2 h_2** | 1.056040534 | 1.53052599 | 188.9792595 | 26.51270145 | 0.23169627 | 47.77805207 | 19.8770936 | 0.80984755 | 319.026459 | 7.16026815 |  |
| **2 h_3** | 1.025365907 | 1.063196975 | 20.67417721 | 1.541182475 | 0.08043194 | 41.44618892 | 18.2038865 | 0.80583157 | 433.753378 | 3.04421442 |  |
| **2 h_4** | 0.089414502 | 0.146739538 | 28.26238182 | 1.549688533 | 0.03195986 | 5.623302291 | 1.23421279 | 0.10065053 | 48.2670426 | 0.57422444 | 0.21660018 |
| **2 h_5** | 0.078434624 | 0.076010474 | 16.43421435 | 0.942469037 | 0.0152514 | 2.825529937 | 0.64114216 | 0.03888084 | 27.6095091 | 0.31388303 | 0.13572061 |
| **2 h_6** | 0.086163843 | 0.135177964 | 21.58536851 | 1.204979417 | 0.02632103 | 5.318585322 | 0.74442375 | 0.05243417 | 34.2503165 | 0.38984257 | 0.18491371 |
| **4 h_1** |  |  |  |  |  |  |  |  |  |  |  |
| **4 h_2** | 0.952244772 | 1.286994219 | 153.7329096 | 24.83423911 | 0.27367595 | 47.67942111 | 25.6093586 | 0.83326314 | 344.078166 | 7.37063509 |  |
| **4 h_3** | 0.860667505 | 1.021593341 | 56.86769599 | 5.429883728 | 0.15236617 | 34.6274072 | 25.9385764 | 0.837183 | 363.185891 | 4.2553842 |  |
| **4 h_4** | 0.077760514 | 0.094733684 | 16.80646924 | 0.985730061 | 0.021761 | 3.554062913 | 1.22538145 | 0.05308938 | 25.8253358 | 0.38231228 | 0.1003137 |
| **4 h_5** | 0.106403476 | 0.142524551 | 36.45205645 | 1.988305175 | 0.06415652 | 4.657450013 | 2.24954817 | 0.08589315 | 48.541091 | 0.78945153 | 0.23856043 |
| **4 h_6** | 0.067299839 | 0.093956226 | 14.38904379 | 0.962417095 | 0.03062466 | 3.42710558 | 1.07437076 | 0.07011953 | 20.0651033 | 0.45309675 | 0.10903428 |
| **6 h_1** |  | 1.337447092 | 11.99688107 | 1.74708967 |  | 34.18664293 | 20.2782417 | 2.34073117 | 605.219785 | 2.06071084 |  |
| **6 h_2** | 0.743094315 | 1.022562573 | 158.121151 | 31.64373514 | 0.26729271 | 38.18131164 | 24.7673972 | 0.97436702 | 252.191775 | 8.47610658 |  |
| **6 h_3** | 0.741888256 | 0.760992577 | 59.65754799 | 7.69568586 | 0.15249779 | 23.39441147 | 17.7516597 | 0.75121892 | 243.49534 | 3.45541105 |  |
| **6 h_4** | 0.049809581 | 0.091425684 | 15.04084536 | 1.006521907 | 0.01934744 | 3.010610437 | 0.78702291 | 0.04269812 | 18.2384419 | 0.36924018 | 0.12707087 |
| **6 h_5** | 0.070987981 | 0.132177945 | 28.81395561 | 1.725399898 | 0.03352028 | 3.397813393 | 1.33335987 | 0.06190763 | 30.5926377 | 0.61501402 | 0.20085269 |
| **6 h_6** | 0.046289866 | 0.067571916 | 8.926150131 | 0.569338667 | 0.01749262 | 1.82289292 | 0.34722313 | 0.04185043 | 9.62185441 | 0.20053034 | 0.09533555 |
| **24 h_1** |  | 1.220537966 | 33.29620818 | 3.962849098 |  | 33.99229108 | 30.9476146 | 3.83787356 | 657.777795 | 4.41659678 |  |
| **24 h_2** | 0.629160814 | 0.738002039 | 112.4907222 | 21.92142108 | 0.30590268 | 35.99950612 | 17.691173 | 0.91748175 | 204.524558 | 6.34833252 |  |
| **24 h_3** | 0.687773046 | 0.805972575 | 93.50726422 | 13.58971417 | 0.2458873 | 24.0072565 | 24.968994 | 1.21352598 | 291.428653 | 5.60460802 |  |
| **24 h_4** | 0.06120579 | 0.112989548 | 25.08262848 | 1.988193905 | 0.05630189 | 4.492998981 | 1.48735476 | 0.08864149 | 30.6986393 | 0.78581047 | 0.36367082 |
| **24 h_5** | 0.032534334 | 0.071296128 | 8.177845535 | 0.740018001 | 0.01660452 | 1.812999241 | 0.39072898 | 0.0878533 | 9.93410607 | 0.30474998 | 0.12564267 |
| **24 h_6** | 0.061380461 | 0.127959773 | 30.47446518 | 2.562543528 | 0.06568598 | 4.440443488 | 1.01809013 | 0.10508835 | 35.7251538 | 1.0201991 | 0.5985944 |
| **48 h_1** |  |  |  |  |  |  |  |  |  |  |  |
| **48 h_2** | 0.681761424 | 0.603237206 | 75.64526093 | 13.98787112 | 0.28194681 | 29.39692984 | 9.09920945 | 1.39837692 | 170.349296 | 4.35403274 |  |
| **48 h_3** | 0.749518708 | 0.875127895 | 165.0883635 | 18.43543643 | 0.53567632 | 32.02165149 | 29.7498713 | 1.7029747 | 560.322616 | 7.79296106 |  |
| **48 h_4** | 0.061536773 | 0.108048874 | 23.26122421 | 1.995036867 | 0.02962256 | 5.624638296 | 3.21948992 | 0.09630936 | 39.4916001 | 0.70909885 | 0.10929763 |
| **48 h_5** | 0.078036846 | 0.113790855 | 32.32729576 | 2.601277782 | 0.02336041 | 8.680408444 | 4.32736251 | 0.13769877 | 46.2679174 | 1.06743222 | 0.11574465 |
| **48 h_6** | 0.080761793 | 0.095769324 | 29.89527923 | 2.672736937 | 0.02152871 | 6.521220453 | 4.50410418 | 0.11461068 | 40.1320361 | 1.20807787 | 0.07705603 |

Replicates 1 to 3 and 4 to 6 were measured in the same experiment, respectively. Data was normalized to control (100 %) for comparison. 1 to 3 was normalized to respective control, 4 to 6 were normalized to control_4. Not every metabolite was detected in every replicate and respective values were left blank. For replicate 4 to 6, 2 technical replicates were performed, here the mean value is depicted.

**Supplemental Table S2. Summary of quantification method for ^15^N-labeled amino acids measured via LC-qTOF.** Shown are all theoretically possible isotopomers after ^15^N-labelling. Isotopomer masses colored in red could not be determined in any sample due to sensitivity or absence in the sample. Natural abundances were retrieved from MassHunter Isotope Distribution Calculator version B7024.29.

|  |  |  | **Mass of isotopomer (m/z)** | | | | | **Natural abundance of isotopomer (%)** | | | | |
| --- | --- | --- | --- | --- | --- | --- | --- | --- | --- | --- | --- | --- |
| **Compound** | **Formula** | **Retention time (min)** | **m0** | **m1** | **m2** | **m3** | **m4** | **m0** | **m1** | **m2** | **m3** | **m4** |
| alpha-Alanine | C3H7NO2 | 2.92 | 90.05495 | 91.05779 |  |  |  | 95.91521 | 3.62391 |  |  |  |
| Arginine | C6H14N4O2 | 20 | 175.11895 | 176.12125 | 177.12328 | 178.12547 | 179.12755 | 91.7848 | 7.52586 | 0.65109 | 0.03661 | 0.00158 |
| Asparagine | C4H8N2O3 | 3.39 | 133.06077 | 134.06332 | 135.06512 |  |  | 94.30294 | 4.97424 | 0.68922 |  |  |
| Aspartate | C4H7NO4 | 2.2 | 134.04478 | 135.04775 |  |  |  | 94.42833 | 4.66098 |  |  |  |
| Cysteine | C3H7NO2S | 2.58 | 122.02703 | 123.02926 |  |  |  | 91.10993 | 4.16172 |  |  |  |
| Glutamate | C5H9NO4 | 2.66 | 148.06043 | 149.06348 |  |  |  | 93.39646 | 5.64169 |  |  |  |
| Glutamine | C5H10N2O3 | 3.63 | 147.07642 | 148.07912 | 149.08089 |  |  | 93.27244 | 5.95014 | 0.73626 |  |  |
| Glycine | C2H5NO2 | 2.5 | 76.0393 | 77.0419 |  |  |  | 96.97491 | 2.59279 |  |  |  |
| Histidine | C6H9N3O2 | 14.36 | 156.07675 | 157.07927 | 158.08125 | 159.08356 |  | 92.1731 | 7.16796 | 0.62333 | 0.03412 |  |
| Isoleucine | C6H13NO2 | 6.2 | 132.10191 | 133.10501 |  |  |  | 92.8051 | 6.58171 |  |  |  |
| Leucine | C6H13NO2 | 5.72 | 132.10191 | 133.10501 |  |  |  | 92.8051 | 6.58171 |  |  |  |
| Lysine | C6H14N2O2 | 11.37 | 147.1128 | 148.11562 | 149.11753 |  |  | 92.45666 | 6.9054 | 0.60417 |  |  |
| Methionine | C5H11NO2S | 4.2 | 150.05833 | 151.06095 |  |  |  | 89.12962 | 6.04027 |  |  |  |
| Phenylalanine | C9H11NO2 | 6.56 | 166.08626 | 167.08943 |  |  |  | 89.87849 | 9.26979 |  |  |  |
| Proline | C5H9NO2 | 2.88 | 116.0706 | 117.07364 |  |  |  | 93.85202 | 5.5977 |  |  |  |
| Serine | C3H7NO3 | 2.32 | 106.04987 | 107.05272 |  |  |  | 95.68214 | 3.65155 |  |  |  |
| Threonine | C4H9NO3 | 2.6 | 120.06552 | 121.06849 |  |  |  | 94.63657 | 4.65698 |  |  |  |
| Tryptophane | C11H12N2O2 | 11.13 | 205.09715 | 206.10019 | 207.10256 |  |  | 87.63511 | 11.26432 | 1.02654 |  |  |
| Tyrosine | C9H11NO3 | 4.2 | 182.08117 | 183.08435 |  |  |  | 89.66009 | 9.28141 |  |  |  |
| Valine | C5H11NO2 | 4 | 118.08626 | 119.0893 |  |  |  | 93.83044 | 5.618 |  |  |  |

**Supplemental Table S3. Raw data from quantification method for ^15^N-labeled amino acids measured via LC-qTOF.**

| **rel. enrichment m+1** | **alpha-Alanine** | **Aspartate** | **Glutamate** | **Histidine** | **Isoleucine** | **Leucine** | **Proline** | **Valine** |
| --- | --- | --- | --- | --- | --- | --- | --- | --- |
| **control_1** | 0 | 0.40090837 | 0.47125559 | 0.91527038 | 0 | 0.05577544 | 0 | 0.20031903 |
| **control_2** | 0 | 0.48510137 | 1.19260657 | 0.37448333 | 0 | 0.80310617 | 0.34308631 | 0 |
| **control_3** |  |  | 1.29267167 | 0 | 0 | 0 | 0 | 0 |
| **1 h_1** | 1.04923406 | 7.42071568 | 7.30439269 | 2.4884458 | 1.10860703 | 1.10742484 | 3.82161783 | 0 |
| **1 h_2** | 1.1342786 | 5.57061267 | 8.08730538 | 1.33380341 | 0.5122767 | 0.16596954 | 4.38318515 | 0.54090537 |
| **1 h_3** | 1.4235847 | 11.7719037 | 7.05797965 | 2.94279624 | 1.56580823 | 1.87013479 | 3.25527612 | 0 |
| **2 h_1** | 1.02353373 | 4.94454043 | 10.6995843 | 0 | 1.6910971 | 2.21479582 | 7.38352294 | 0.94699479 |
| **2 h_2** | 1.50847976 | 11.6258644 | 9.14597736 | 0.15150108 | 2.01387955 | 4.44076122 | 6.13227587 | 0.77000518 |
| **2 h_3** | 1.63380619 | 7.96397967 | 10.0765488 | 3.40861241 | 1.21635719 | 1.44060611 | 6.73474321 | 0.31476989 |
| **4 h_1** | 2.77573225 | 17.8409345 | 22.2289823 | 0.73633046 | 2.64423697 | 4.70825532 | 15.2264821 | 1.21290403 |
| **4 h_2** | 1.58387165 | 15.1567539 | 19.8113117 | 0.27736634 | 2.20542332 | 4.1037447 | 13.5017598 | 1.15444201 |
| **4 h_3** | 3.25053851 | 17.4705266 | 22.8746761 | 0 | 3.05079323 | 5.74084546 | 15.1134947 | 1.43422227 |
| **6 h_1** | 1.48273715 | 11.1146972 | 15.724708 | 0.07174415 | 2.13343836 | 3.26234773 | 12.6294388 | 1.31192073 |
| **6 h_2** | 1.04540805 | 10.6964563 | 14.9846617 | 0.40007852 | 2.37162829 | 4.13902977 | 11.8173998 | 1.51042917 |
| **6 h_3** | 1.81965465 | 15.2847053 | 18.0640338 | 0 | 3.12946529 | 4.56325965 | 14.8371646 | 1.8754336 |
| **24 h_1** | 1.91397889 | 15.4363329 | 20.3238226 | 0.52366814 | 2.48802449 | 3.60320655 | 18.6531695 | 1.21526608 |
| **24 h_2** | 2.22740472 | 16.250518 | 20.3090225 | 0 | 2.99205761 | 5.24440308 | 18.3253367 | 1.25429893 |
| **24 h_3** | 2.47775372 | 16.5117326 | 18.2038927 | 1.00092673 | 3.74397819 | 5.57323623 | 17.2402266 | 2.17686156 |
| **48 h_1** | 1.50324848 | 8.96337872 | 12.0635386 | 0.29616694 | 2.46188169 | 3.6951194 | 11.187649 | 0.76294907 |
| **48 h_2** | 1.31638932 | 9.8710555 | 13.2015612 | 0 | 2.05882482 | 3.43128094 | 12.2095382 | 0.93567938 |
| **48 h_3** | 1.2572175 | 10.4702217 | 14.2606943 | 0.48794577 | 2.39205954 | 3.96856915 | 11.7648981 | 3.40078421 |

Shown are only metabolites with a relative enrichment of ^15^N of at least 1 % at any time point.

**Data analysis for ^15^N-labeled amino acids.** Amino acid peaks in the samples were identified at mass-to-charge (m/z) ratios and retention times (RT) listed above and with external amino acid standards measured in parallel. Peaks were integrated via Agilent Mass Hunter Workstation B07 (Agilent Technologies, Santa Clara CA, USA). Peak intensities of m1 (i.e. incorporation of one ^15^N-label) were corrected for their natural abundance via the following calculation.

m0_int_, peak intensity of non-labelled amino acid; m1_int_, peak intensity of amino acid with one ^15^N-label (m+1); m1_nat_, natural abundance of isotopomer (see above);

m1_background_	= m0_int_ • m1_nat_ : 100;	m1_enrichment_	= m1_int_ - m1_background;_	m1_relative.enrichment_	= m1_enrichment_ : m0_int_ • 100
